# Brain Diffusion Transformer for Personalized Neuroscience and Psychiatry

**DOI:** 10.1101/2025.04.12.648506

**Authors:** Rongquan Zhai, Yechen Hu, Liping Zheng, Shitong Xiang, Chao Xie, Lei Peng, Tobias Banaschewski, Gareth J. Barker, Arun L.W. Bokde, Rüdiger Brühl, Sylvane Desrivières, Herta Flor, Hugh Garavan, Penny Gowland, Antoine Grigis, Andreas Heinz, Herve Lemaitre, Jean-Luc Martinot, Marie-Laure Paillère Martinot, Eric Artiges, Frauke Nees, Dimitri Papadopoulos Orfanos, Luise Poustka, Michael N. Smolka, Sarah Hohmann, Nathalie Holz, Nilakshi Vaidya, Robert Whelan, Zuo Zhang, Lauren Robinson, Jeanne Winterer, Sinead King, Yuning Zhang, Hedi Kebir, Ulrike Schmidt, Julia Sinclair, Argyris Stringaris, Gunter Schumann, Danilo Bzdok, Henrik Walter, Edmund T. Rolls, Barbara Sahakian, Trevor W. Robbins, Jianfeng Feng, Weikang Gong, Tianye Jia, IMAGEN Consortium, STRATIFY Consortium

## Abstract

Task-fMRI analyses typically focus on localized activation contrasts between stimuli, neglecting the brain’s dynamic hierarchy. We introduce Brain Diffusion Transformer (Brain-DiT), a deep generative model capturing recurrent processing underlying individualized neurocognitive state transitions via functional networks. Without prior assumptions, Brain-DiT identifies canonical cognitive regions in the brain and reveals replicable subgroups with distinct neural circuits in large cohorts, offering critical clinical insights overlooked by traditional methods: individuals exhibiting negative emotion bias, linked to language-related regions, had a 12-fold higher likelihood of major depression, and those with maladaptive inhibition strategies, associated with overactive medial frontal regions, showed a 9-fold increased risk of alcohol abuse. By bridging cognitive theory and psychiatric applications, Brain-DiT provides a unified analytical paradigm, paving the way for operational personalized medicine in psychiatry.

## Introduction

Brain activation and functional connectivity are central to fMRI studies, offering essential and complementary insights into brain function (*1, 2*). Together, brain activation and functional connectivity provide a high-dimensional dynamic overview of how signals are transferred and communicated across the brain’s highway along the neural processing hierarchy. While event-evoked brain activation represents the traffic formed by various types of vehicles (*i.e*., the brain’s cognitive state), the functional connectivity network reflects the highway’s status or condition that determines how swiftly and efficiently the traffic can move (*3*). Previous studies have attempted to understand the relationship between these two distinct measures, such as using functional connectivity to predict localized brain activation (*4, 5*). However, few have integrated these measures into a unified and comprehensive model that is sophisticated enough for a high-dimensional, recurrent, and dynamic system like the human brain. Here, we propose a novel analytical framework that combines localized brain activation and brain-wide communication to systematically reveal how high-dimensional dynamic neural networks support personalized adaptive cognition, a hallmark of brain function (*6*).

The aim of our present study is to advance the analysis of neuroimaging data by producing a generative model that can predict task activation in a second condition (such as viewing a neutral face) from the neuroimaging activation from a first task condition (such as viewing an angry face) by using additional information such as the functional connectivity for each individual (**Fig. 1A**). Particularly, we utilize the state-of-the-art diffusion generative model (*7-14*) to model all plausible transition neuropathways between cognitive states for different task conditions, instead of a traditional direct contrast of activation (*e.g*., angry > neutral) (**Fig. 1B**). The deployment of the diffusion generative model is not only because of its superior robustness for both training and application compared to other mainstream neural network models (*15*), but also because its recursive signal updating, when guided by functional connectivity constraints, parallels the brain’s hierarchical processing of information (*16*). Notably, a direct connection between the hierarchical generative model and the canonical microcircuit has long been proposed (*17*). We show that the understanding developed in the generative model helps to interpret neuroimaging data related to depression and alcohol abuse. Particularly, this new approach enables the prediction of individualized distinct characteristics solely based on neuroimaging features, such as biased attention to negative stimuli (leading to depression) or maladaptive inhibition strategies (underlying alcohol abuse). In independent clinical data, the predictions from the generative model are much better (by 5-fold) than when the prediction is based just on the activation and/or functional connectivity.

**Fig. 1.**
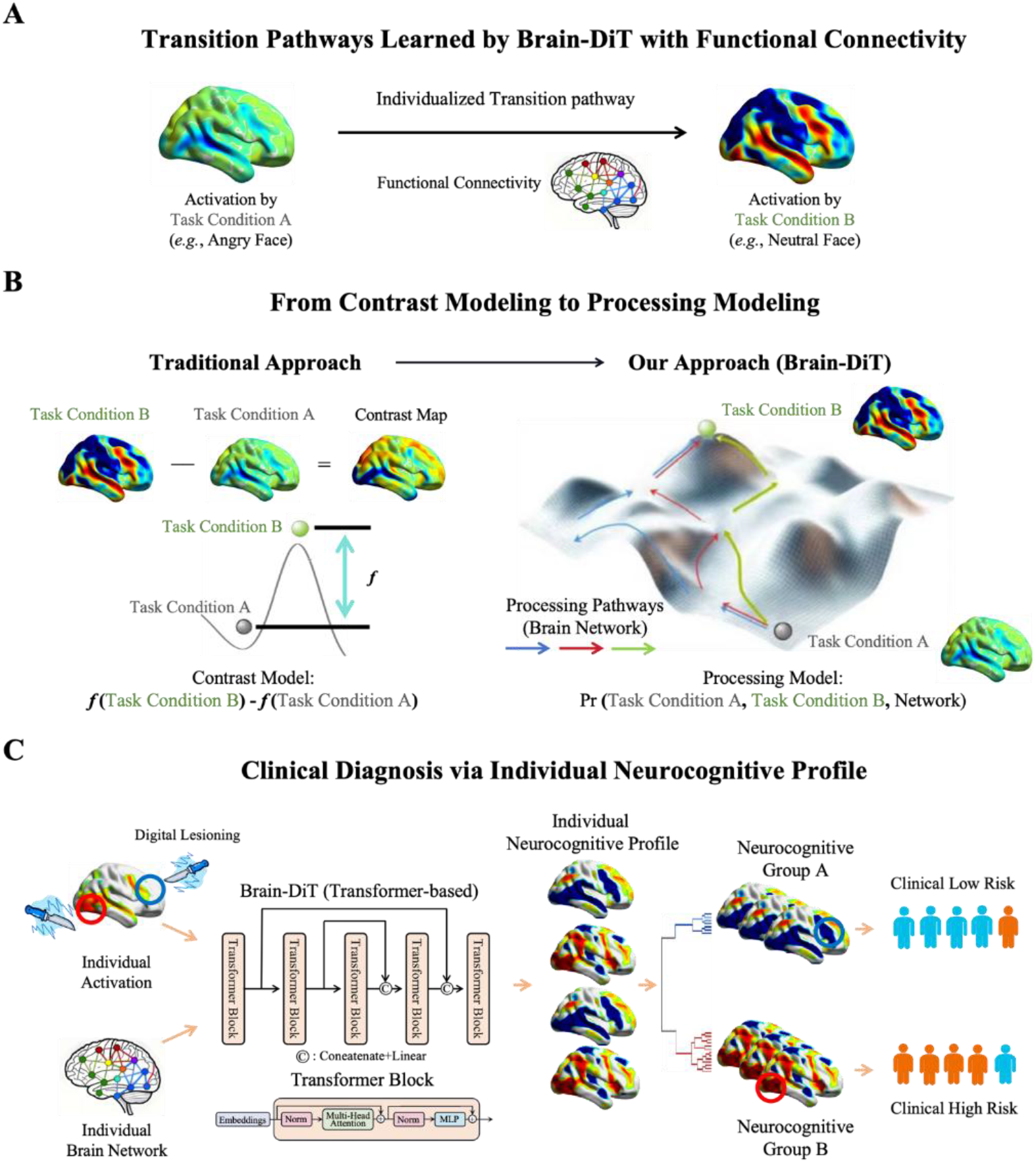
Overview of Neurocognitive Processing Modeling and Brain Diffusion Transformer (Brain-DiT). **A**. Brain-DiT is a generative model to learn how brain activation to one stimulus can be used to predict brain activation to a different stimulus through functional connectivity. **B**. Compared to the traditional contrast model, Brain-DiT models the joint probability of brain activation and brain networks during transitions between cognitive states under different task conditions. **C**. By utilizing the established Brain-DiT model, digital lesioning enables us to obtain individualized neurocognitive profiles that summarize each brain region’s importance to the Brain-DiT model, thus allowing classifications based on individualized processing pathways. These neuroimaging-based classifications exhibit remarkably high agreement with clinical disease categorizations.

## Results

### A Diffusion Transformer Model Tailored to Neuroimaging

Based on a diffusion transformer model widely used in the field of image generation (*18*), we establish Brain Diffusion Transformer (Brain-DiT). The model operates by taking an individual’s task-related brain activation map (such as a response to an angry face) as the source condition, along with this participant’s resting-state functional connectivity network. It then predicts a target activation pattern associated with a related but different task condition (such as a neutral face) (**Fig. 1A & fig. S1**). The detailed operation of Brain-DiT is described in the Supplementary Material. A key advantage of Brain-DiT is that it learns relations between the two critical measures of brain function, *i.e*., a joint function of activation in a brain region and functional connectivity between brain regions (**Fig. 1B**). Moreover, the learned Brain-DiT model serves as a powerful analytical tool for assessing the brain of each individual, such as identifying a tendency for depression, or for alcohol abuse. To make an inference about differences between people, we applied an approach to removing parts of the activation map (termed digital lesioning; **Fig. 1C**) in order to find which brain regions were important in the mapping learned by Brain-DiT. This cognitive relevance map for each individual was used to cluster participants (**Fig. 1C**) for further behavioral and clinical characterizations.

To demonstrate the model’s broad applicability, we trained Brain-DiT on fMRI images from two distinct tasks in the IMAGEN dataset (*19*): the block-design passive viewing emotional face task (EFT), which assesses emotional processing, and the event-design proactive stop-signal task (SST), which is a widely accepted task to examine inhibitory control. These data were collected at three time points: age 14 (baseline, BL), age 19 (follow-up 2, FU2), and age 23 (follow-up 3, FU3). Each longitudinal data point was treated as an independent data entry in the model. Brain-DiT was trained and validated with fine-tuned hyperparameters on EFT (*n*_training_ = 2720 and *n*_validation_ = 302) and SST (*n*_training_ = 2412 and *n*_validation_ = 268), then tested on independent hold-out datasets (*n* = 689 for EFT and *n* = 729 for SST). To minimize the influence of potential confounding factors, we carefully matched the training, validation, and testing samples by gender, research sites, and longitudinal information (see details in Supplementary Materials).

In the testing sample, Brain-DiT achieved remarkable individualized (**Fig. 2 A&C**) brain-wide prediction accuracy (or transformation performance) for the target brain activation (EFT neutral face: Pearson’s *r*_mean_similarity_ = 0.74, *P*_perm_10000_ = 5.0×10^−4^ and SST stop-success *r*_mean_similarity_ = 0.72, *P*_perm_10000_ = 9.0×10^−4^; *r* represents Pearson’s correlation throughout the manuscript; see more numerical evaluations in **Fig. 2 B&D** and **fig. S2**&**S3**). Both brain activation and functional connectivity were essential for accurate predictions, as a model trained without either input could only predict random outcomes (*r*_mean_similarity_ < 0.01, *P*_perm_10000_ > 0.79 for both EFT and SST; **fig. S4 A&D**). Also, Brain-DiT outperformed alternative models (*i.e*., with different MRI modalities or linear models (*20*); see **fig. S4 B, C, E&F**). Finally, cross-task predictions (*i.e*., from EFT to SST and vice versa) failed to achieve satisfactory accuracy (*r*_mean_similarity_ < 0.40, *P*_perm_10000_ > 0.37, **fig. S5**), reinforcing that EFT and SST rely on distinct cognitive processes (**fig. S6**).

**Fig. 2.**
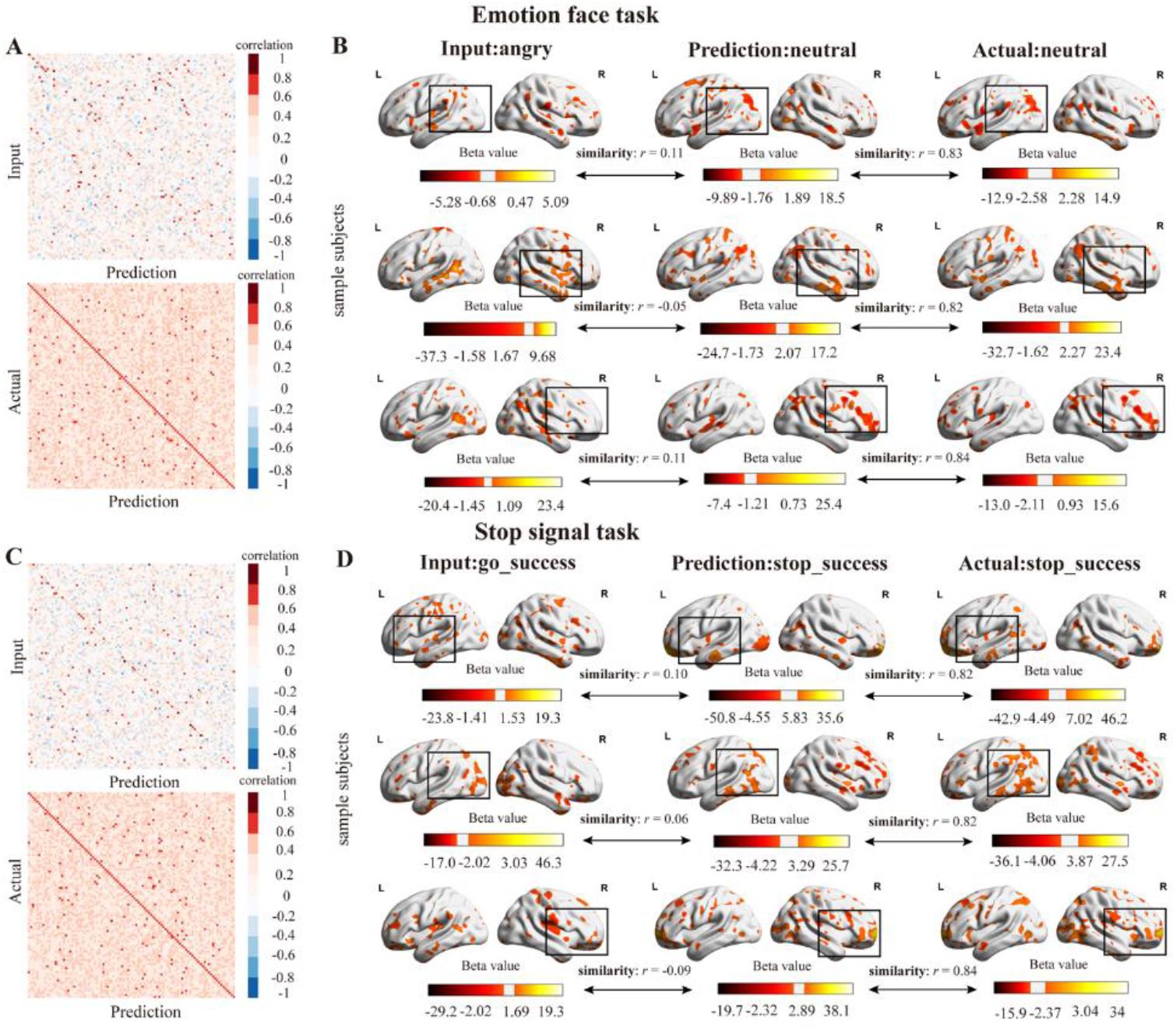
Individual generation performance. **A**. The input-actual and prediction-actual correlation matrices of the emotion face task for 100 random samples in the test data. **B**. In sample individuals across the three timepoints, the predicted neutral face activation maps resemble the actual activation (*r* >= 0.82), while being distant from the input angry face condition activation (|*r*| <= 0.11). **C**. The input-actual and prediction-actual correlation matrices of the stop-signal task for 100 random samples in the test data. **D**. In sample individuals across the three timepoints, the predicted stop-success face activation maps resemble the actual activation (*r* >= 0.82), while being distant from the input go-success condition activation (|*r*| <= 0.10).

## Digital Lesioning with Brain-DiT

We then performed digital lesioning with the established Brain-DiT (**Fig. 1C**) to investigate the causal influence of each brain region on the model’s performance. In detail, by systematically inactivating part of the brain—setting its activation to zero—we performed inference in the testing sample using the established generative model, which was trained with pre-lesion brain activation, to assess the impact on model predictions. First, we inactivated a portion of the brain, as indicated by the meta-analysis activation maps (roughly 50% of the brain), for the cognitive domains ‘emotional’ and ‘inhibition’, as derived from the Neurosynth database (*21*) (**Fig. 3A, G**). This lesion process caused the model’s performance to collapse (EFT: *r*_mean_similarity_ dropped from 0.74 to 0.13, *P*_perm_10000_ = 0.31; SST: *r*_mean_similarity_ dropped from 0.72 to 0.08, *P*_perm_10000_ = 0.75). In contrast, when using only the respective meta-maps as input, Brain-DiT maintained much of its predictive performance (*r*_mean_similarity_ = 0.52, *P*_perm_10000_ = 3.3×10^−3^ for EFT; *r*_mean_similarity_ = 0.56, *P*_perm_10000_ = 2.4×10^−3^ for SST; **Fig. 3B, H**). These findings suggest that Brain-DiT effectively identified the necessity and sufficiency of critical brain regions in cognitive processing—without prior knowledge.

**Fig. 3.**
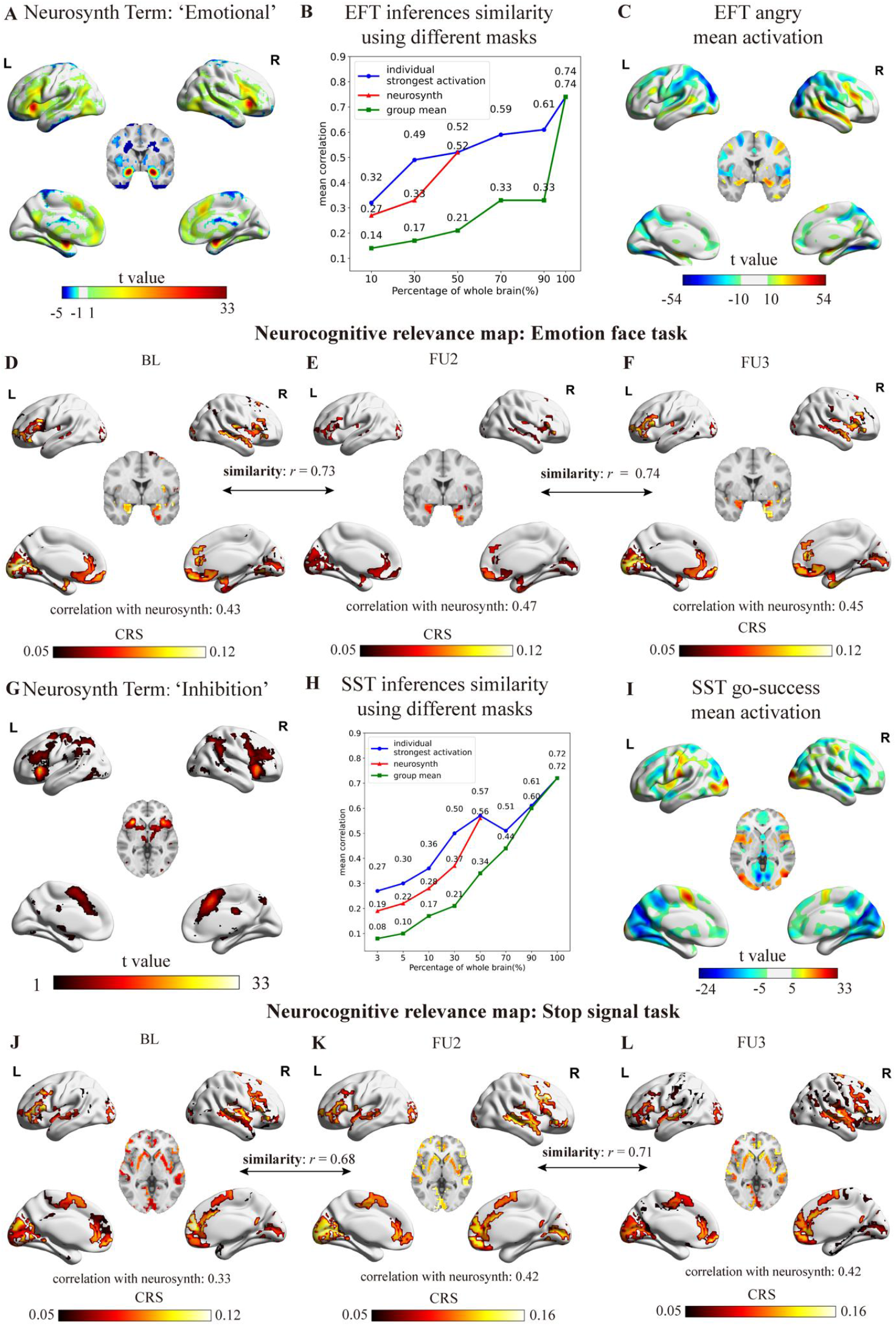
Evaluation of cognitive sensitivity of generative cognitive diffusion transformers (Brain-DiTs) through digital lesioning experiments and cognitive relevance mapping. **A**. Term “Emotional” meta map from Neurosynth(*21*) database. **B**. The mean prediction accuracy (*i.e*., the spatial similarity between predicted and actual activation maps) for the neutral face condition from established Brain-DiT, using different proportions of masks (*i.e*., the individualized angry face activation map, the “emotional” meta map from Neurosynth(*21*) and the group mean of angry face activation map). **C**. The group mean activation map of the angry face condition in the training sample. **D-F**. The cognitive relevance score maps of the emotional face task, acquired at BL, FU2, and FU3. **G**. Term “Inhibition” meta map from Neurosynth(*21*) database. **H**. The mean prediction accuracy for the go-success condition from established Brain-DiT, using different proportions of masks (*i.e*., the individualized stop-success activation map, the “Inhibition” meta map from Neurosynth(*21*) and the group mean of stop-success activation map). **I**. The mean activation map of the stop-success condition in the training sample. **J-L**. The cognitive relevance score maps at BL, FU2, and FU3 of the stop-signal task. Abbreviations: BL (baseline), FU2 (follow-up 2), and FU3 (follow-up 3).

Next, we systematically lesioned the brain, preserving 90%, 70%, 50%, 30%, or 10% (from high to low based on significance) of each participant’s brain through three different strategies: (1) the individual activation map (I-map), (2) the Neurosynth meta map (N-map, starting from 50% of the brain), and (3) the group-level mean activation map (G-map). For both EFT and SST, I-map was the most robust, while G-map retained the least transformation performance, with N-map in between (*t*_EFT_I>N_ = 3.30 and *t*_EFT_N>G_ = 3.02, both *P*_*FDR*_ < 0.010; *t*_SST_I>N_ = 2.85 and *t*_SST_N>G_ = 2.75, both *P*_*FDR*_ < 0.010; the false discovery rate (*FDR*) adjusted for the two comparisons in EFT and SST, respectively; **Fig. 3 B&H**). These results indicate that both shared core regions and personalized features are critical for adaptive cognitive processing.

Notably, in EFT, a substantial gap (*r*_similarity*_*diff_ = 0.28; **Fig. 3B**) was observed between the I-map and G-map even after the least important 10% of the brain was lesioned, suggesting a very high degree of heterogeneity in emotional processing (*22, 23*), where distinct but minor personalized cognitive features (for instance, subconscious perception of emotional signals) may play a vital role. This phenomenon was not observed for the SST (**Fig. 3H**), hence suggesting more homogenous cognitive processing in motor inhibition (*24, 25*).

Finally, based on the HCPex template (*26*), we sequentially lesioned each brain region and assigned the decrease in transformation performance as a measurement of this region’s cognitive importance. This process yields individualized brain-wide cognitive relevance score (CRS) maps for both EFT and SST. The population mean CRS maps were highly consistent across all data timepoints (*r* >= 0.73, *P*_perm_10000_ <= 4.0×10^−4^ for EFT and *r* >= 0.68, *P*_perm_10000_ <= 5.0×10^−4^ for SST) (**Fig. 3D-F, J-L**), reassuring Brain-DiT’s ability to reliably identify critical brain regions involved in cognitive processing.

### Qualitatively Stratified Neurocognitive Profiles

At each data collection timepoint (ages 14, 19, and 23) of the IMAGEN project, we applied hierarchical clustering on individualized CRS maps to identify subgroups with distinct neural circuit engagement (or neurocognitive profiles). The silhouette coefficient overwhelmingly suggested a two-cluster optimal solution for both EFT and SST across all three timepoints (**table S1**). Remarkably, the brain features that differentiated the stratified subgroups were almost identical over time for both EFT (*r*_similarity_ >= 0.82, *P*_perm_10000_ <= 2.0×10^−4^) and SST (*r*_similarity_ >= 0.84, *P*_perm_10000_ <= 1.0×10^−4^; **Fig. 4 A-C, H-J**), suggesting consistent and qualitative between-group differences in their cognitive predispositions, likely involving implicit preparatory perceptual or response sets (*27*). For both EFT and SST subgroups at each of the three timepoints, we observed no difference (even nominally) in gender and research sites (**table S5**).

**Fig. 4.**
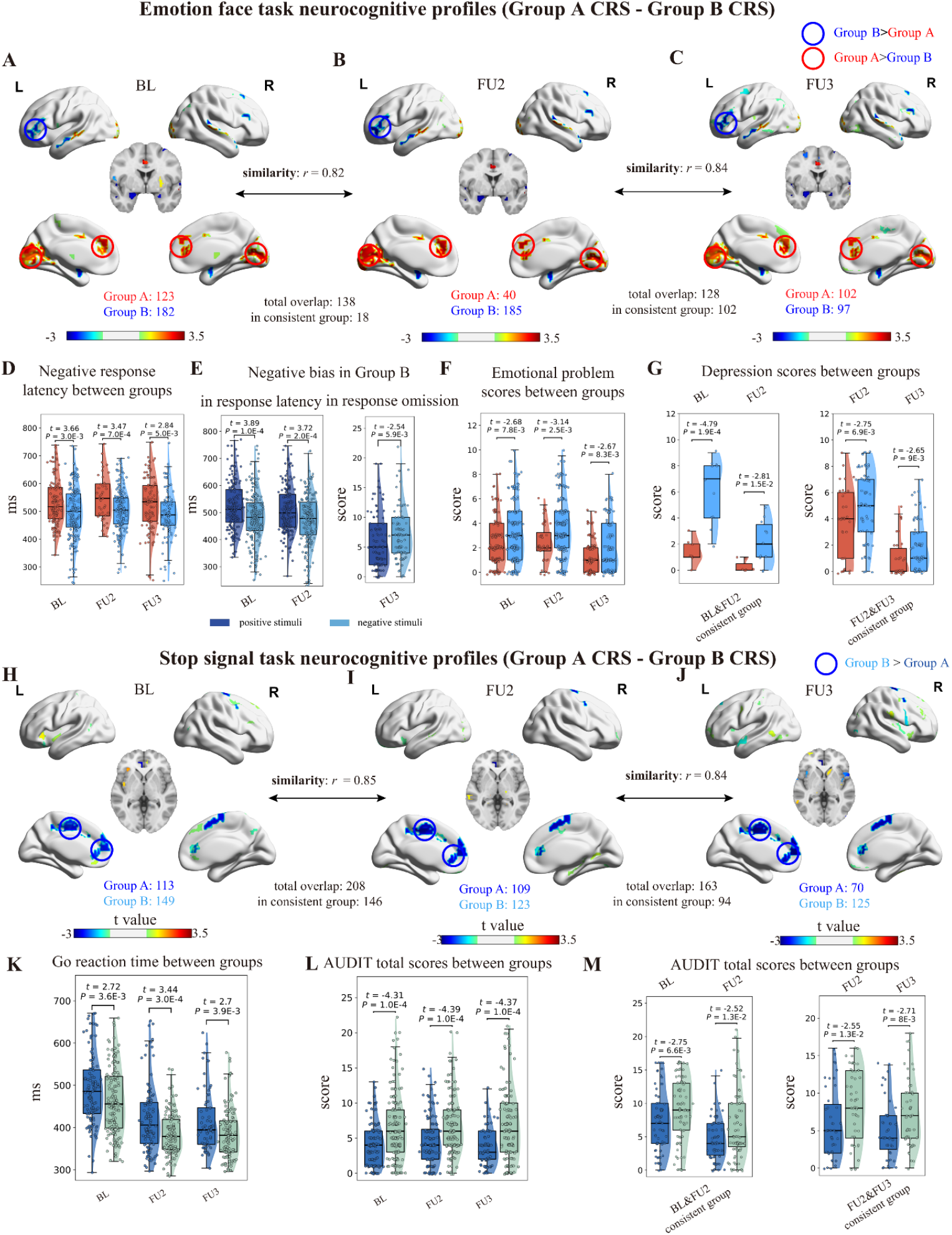
Replicable neurocognitive profiles and the clinical implications in psychiatric disorders. **A-C**. The t-maps highlight the longitudinally consistent neurocognitive profiles that distinguish subgroups in the emotional face task (EFT) (*i.e*., the left inferior frontal cortex for Group B vs the dorsal anterior cingulate cortex, and primary visual cortex for Group A) based on hierarchical clustering at BL (baseline), FU2 (follow-up 2), and FU3 (follow-up 3) respectively. Notably, despite the almost identical neurocognitive profiles of subgroups, very few participants remained in the same subgroups from BL to FU2, while most participants retained the same subgroups from FU2 to FU3. **D-G**. In the IMAGEN dataset, the two EFT subgroups also exhibited longitudinally consistent differences across various emotion-related behaviors. **H-J**. The t-maps highlight the longitudinally consistent subgroup profiles that distinguish subgroups in the stop-signal task (SST) (*i.e*., Group A and Group B were differentially engaged with the pregenual anterior cingulate cortex, medial prefrontal cortex, and supplementary motor area) based on hierarchical clustering at BL, FU2, and FU3 respectively. Unlike the EFT, participants showed high longitudinal consistency in SST subgroups from BL to FU3. **K-M**. In the IMAGEN dataset, the two SST subgroups also exhibited longitudinally consistent differences in alcohol abuse-related behaviors.

### Emotional Processing (EFT Subgroups)

The two subgroups were featured with the greater engagement of the left inferior frontal cortex (left-IFC) (*n* = 182, 185, and 97 at BL, FU2, and FU3) and greater engagement of the dorsal anterior cingulate cortex (dACC) and the primary visual cortex (V1) (*n* = 123, 40, and 102 at BL, FU2 and FU3), respectively. Notably, none of these regions were highlighted in the Neurosynth ‘emotional’ meta-map, thus in line with the digital lesion finding (**Fig. 3D-F**) that heterogeneous perceptions may precede primary emotional processing. Remarkably, despite the longitudinal consistency of the brain features that differentiate the subgroups, very few individuals (18/138, 13%, *P* < 1.0×10^−3^ compared to a random 50%) remained in their groups from age 14 (BL) to age 19 (FU2), whereas most individuals remained stable (102/136, 75%, *P* < 1.0×10^−3^) from age 19 (FU2) to age 23 (FU3) (**Fig. 4A-C**; also see **table S2** for details). Such a drastic dynamic shift in stability may represent the maturation of the emotional regulation system (*28*).

Nevertheless, it was always the left-IFC subgroup that demonstrated a much faster response to negative stimuli during a separately conducted affective go/no-go task (*t* <= -2.84, *Cohen’s d* <= -0.40, *P*_*FDR*_ < 0.05 across the three timepoints; *FDR* adjusted for 12 tests: 3 timepoints x 4 measures; **Fig. 4D** and **table S3**). Further, within the left-IFC subgroup, participants also demonstrated a consistent attentional bias toward negative stimuli, although the bias was primarily expressed as shorter latency at BL (*t* = -3.89, *Cohen’s d* = -0.29, *P*_*FDR*_ = 8.1×10^−3^; **Fig. 4E** and **table S3**) and FU2 (*t* = -3.72, *Cohen’s d* = -0.28, *P*_*FDR*_ = 8.1×10^−3^; **Fig. 4E** and **table S3**), but as increased omissions at FU3 (*t* = 2.54, *Cohen’s d* = 0.26, *P*_*FDR*_ = 0.026; **Fig. 4E** and **table S3;** *FDR* adjusted for 6 tests: 3 timepoints x 2 measures). Such a shift could be a maladaptive outcome of self-management for negative bias (*29*). As expected, the left-IFC subgroup also had higher emotional symptoms across all three timepoints (*t* >= 2.67, *Cohen’s d* >= 0.31, *P*_*FDR*_ < 0.05 for emotional symptoms), though only with significantly increased depression scores at FU3 (*t* = 3.75, *Cohen’s d* = 0.54, *P*_*FDR*_ = 1.4×10^−3^; *FDR* adjusted for 6 tests: 3 timepoints x 2 measures; **Fig. 4F** and **table S3**). Importantly, in contrast to the above findings, participants persistently classified in the left-IFC subgroup had much higher depression scores than those remaining in the dACC/V1 subgroup (from BL to FU2 *n* = 11 vs 7: *t* >= 2.81, *Cohen’s d* >= 1.35, *P*_*FDR*_ <= 0.012 for BL and FU2 scores; from FU2 to FU3 *n* = 68 vs 34: *t* >= 2.65, *Cohen’s d* >= 0.56, *P*_*FDR*_ <= 0.012 for FU2 and FU3 scores; *FDR* adjusted for 4 tests: 2 timespans x 2 measures; **Fig. 4G** and **table S3**), hence suggesting prolonged engagement of the left-IFC in emotional processing as a cognitive basis for depressive symptoms. Finally, if we selected high-risk participants (>50% for diagnosis) for major depression (*n* = 17) and healthy controls (*n* = 41) in FU3 (when depression becomes a more prominent concern), the left-IFC subgroup had a 9-fold increased risk (an odds ratio (*OR*) = 9.0, 95%*CI* = [2.2 ∼ 36.6], *P* = 8.3×10^−4^) compared to the dACC/V1 subgroup (accuracy = 70.7%, recall = 82.4%, precision = 50.0%, F1-score = 62.2%)(**table S4**).

Overall, the above results suggest that participants who perceive emotional visual stimuli primarily using the Broca language area are prone to emotional dysregulation. This unilateral finding in the left-IFC was supported by the collective evidence of 14 transcranial magnetic stimulation (TMS) studies for depression (*30*) and also a recent transcranial direct current stimulation (tDCS) finding, where excitatory/inhibitory regulation of left-IFC could causally impair/facilitate emotional regulation (*31*).

### Motor Inhibition (SST Subgroups)

The two SST subgroups were primarily distinguished by low (*n* = 113, 109, and 70 at BL, FU2, and FU3) versus high (*n* = 149, 123, and 125 at BL, FU2, and FU3) engagement of the pregenual anterior cingulate cortex (pgACC), medial prefrontal cortex (mPFC), and supplementary motor area (SMA), all of which were highlighted in the ‘inhibition’ meta-map from Neurosynth. Such a result aligned with our digital lesion findings that the two subgroups may engage in highly similar motor inhibition processes, though with subtle differences in certain core neural circuits. Unlike the EFT, the subgroups in the SST were relatively stable from BL to FU2 (166/208, 80%, *P* < 1.0×10^−3^) and less so from FU2 to FU3 (94/163, 58%, *P* = 0.050) (**Fig. 4H-J**).

Among all the task behavioral measurements, the two subgroups only differed in reaction time during the go-trials (faster in participants with higher pgACC/mPFC/SMA engagement) but not in stop-signal reaction time. This difference persisted across all three timepoints (*t* <= -2.70, *Cohen’s d* <= -0.34, *P*_*FDR*_ <= 0.015; *FDR* adjusted for 6 tests: 3 timepoints x 2 measures; **Fig. 4K** and **table S6**). However, to our surprise, while the pgACC/mPFC/SMA high-engagement subgroup also exhibited much higher alcohol abuse behavior (a typical addictive symptom linked to inhibitory dysfunction (*32*)) than the other subgroup across all timepoints (*t* >= 4.31, *Cohen’s d* >= 0.54, *P*_*FDR*_ <= 2.3×10^−5^ for timepoints; *t* >= 2.52, *Cohen’s d* >= 0.42, *P*_*FDR*_ <= 0.013 for timespans; **Fig. 4L-M** and **table S6**), we observed no group differences in any measures of impulsiveness (all *P*_*uncorrected*_ > 0.05, and see **table S6** for details). Therefore, the shorter latency in go-trials should not be conventionally interpreted as increased impulsivity. Instead, it may reflect a lack of strategic adjustment, as it is common for participants in SST to slow down to increase the likelihood of stopping their ongoing response (*33*). This proposition also aligns well with pgACC’s role in prospective decision-making, where individuals with greater pgACC engagement demonstrated higher motivation to maintain effortful behavior long-term, despite the cost (*34*). Finally, if we further selected high-risk alcohol dependence individuals (AUDIT total >= 8, *n* = 22) and healthy controls (*n* = 40) in FU3, the pgACC/mPFC/SMA high-engagement subgroup had a 12-fold increased risk (*OR* = 11.9, 95%*CI* = [3.3 ∼ 43.0], *P* = 4.2×10^−5^) compared to the other subgroup (accuracy = 75.8%, recall = 81.8%, precision = 62.1%, F1-score = 70.6%)(**table S4**). Notably, the medial frontal brain regions (pgACC, mPFC, and SMA) were all highlighted in the lesion map and TMS targets aim at reducing alcohol addiction behavior (*35*), thus further supporting the proposed regulatory role of pgACC/mPFC/SMA in alcohol abuse.

### Application and Validation of Brain-DiT in a Clinical Dataset

We then applied the established Brain-DiT and digital lesion approach to the EFT and SST fMRI maps from a clinical dataset Stratify (with an average age of 23) to acquire each participant’s CRS map. Subsequently, we applied hierarchical clustering in case/control studies (*36*) for major depressive disorder (MDD) and alcohol use disorder (AUD). Specifically, we used the EFT Brain-DiT for MDD (*n* = 127) versus controls (*n* = 66) and the SST model for AUD (*n* = 119) versus controls (*n* = 61), according to evidence from IMAGEN. Sex and research sites were matched between cases and controls.

Remarkably, the identified subgroups mirrored those from FU3 (also at age 23) in IMAGEN, displaying nearly identical neurocognitive profiles that distinguish the respective subgroups in both EFT and SST (*r*_EFT_ = 0.86, *P*_EFT-perm_10000_ = 1×10^−4^; *r*_SST_ = 0.90, *P*_SST-perm_10000_ < 1×10^−4^; **Fig. 4O** and **C, P&K**; also see **table S7**). Moreover, these entirely imaging-based neurocognitive subgroups demonstrated remarkably high diagnostic predictivity for MDD (*OR* = 11.6, 95%*CI* = [5.7 ∼ 23.4], *χ*^*2*^ = 53.8, *P* = 2.2×10^−13^; accuracy = 77.7%, recall = 78.7%, precision = 86.2%, F1-score = 82.2%; **Fig. 5A&B**) and AUD (*OR* = 9.2, 95%*CI* = [4.5 ∼ 18.7], *χ*^*2*^ = 42.9, *P* = 5.7×10^−11^; accuracy = 76.7%, recall = 80.7%, precision = 83.5%, F1-score = 82.1%; **Fig. 5D&E**). To assess the specificity of these findings, we conducted a counterfactual analysis by applying the EFT model to AUD and the SST model to MDD. The resulting hierarchical clustering yielded only weak group differences for AUD (*OR* = 1.9, 95%*CI* = [1.0 ∼ 3.6], *χ*^*2*^ = 4.25, *P* = 0.039) and MDD (*OR* = 1.9, 95%*CI* = [1.1 ∼ 3.5], *χ*^*2*^ = 4.56, *P* = 0.033; **table S8**). The nominal significances are likely due to comorbid AUD and MDD, thus supporting the cognitive-diagnostic specificity of Brain-DiT’s predictive power.

**Fig. 5.**
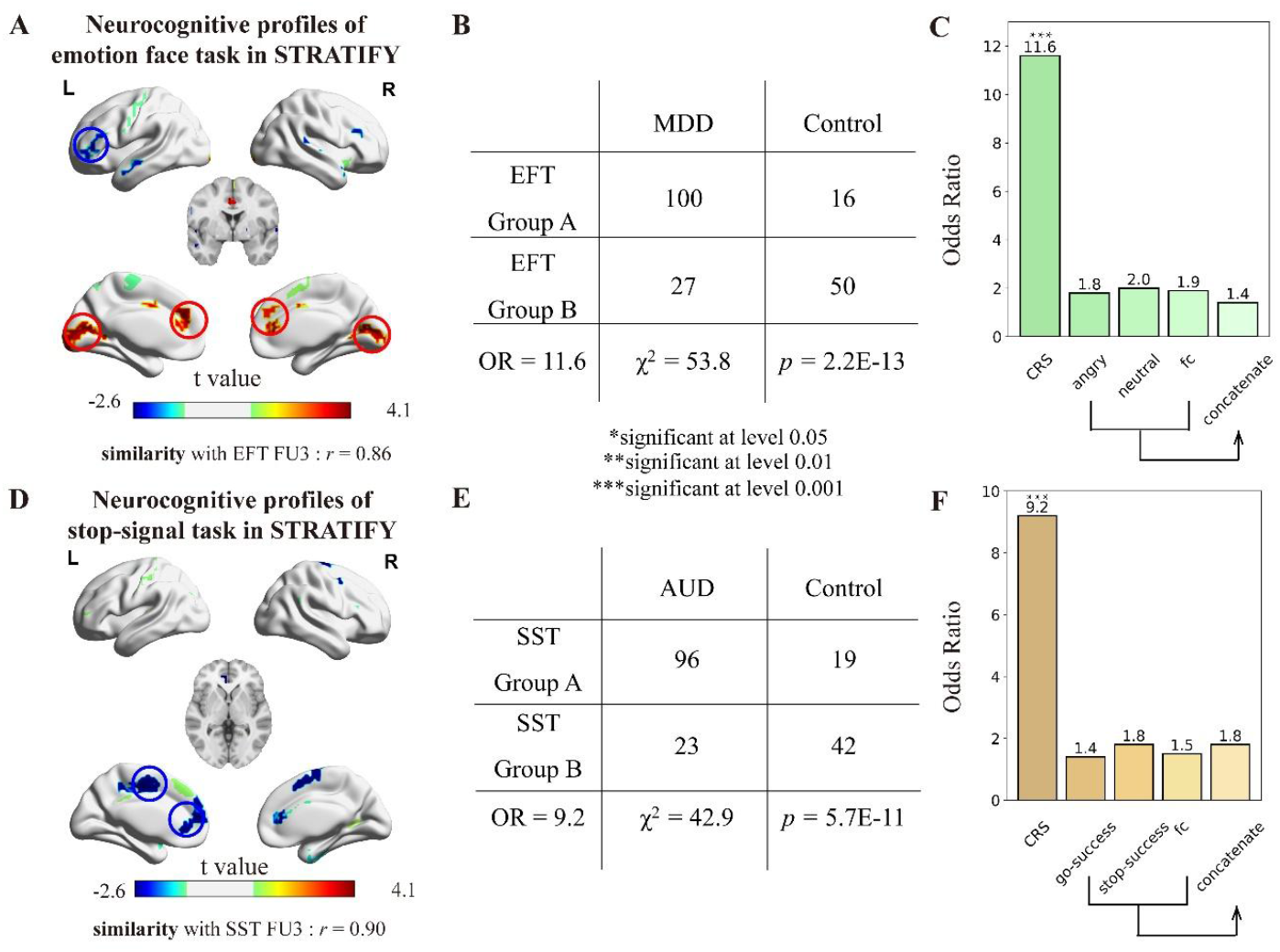
Validation in STRATIFY dataset. In the clinical validation dataset Stratify, almost identical neurocognitive profiles for EFT and SST were again observed between the subgroups derived from hierarchical clustering in the case/control data for major depression disorder (MDD) (**A**) and alcohol use disorder (AUD) (**D**), respectively. Remarkably, these subgroups showed an exceptionally high overlap with the diagnostic groups of MDD (**B**) and AUD (**E**). The cognitive relevance score is the most effective feature for distinguishing between patients and healthy individuals, compared to traditional activation and resting-state functional connectivity (**C&F**).

### Comparison with Traditional Measures

Finally, we demonstrated that traditional brain fMRI measures showed limited ability to discriminate between cases and controls in Stratify using HCPex-defined brain regions. Specifically, hierarchical clustering of individualized brainwise activation or functional connectivity resulted in *OR* <= 2.0 (*P* > 0.05) for MDD (**Fig. 5C**) and *OR* <= 1.8 (*P* > 0.05) for AUD (**Fig. 5F**; also see **tables S9** for more results). Further, we also demonstrated that the CRS acquired from the proposed Brain-DiT could not be derived from a linear combination (the Ridge regression) of activation and functional connectivity (all *P*_Imagen_EFT_ > 0.05, the averaged coefficient of determinant of multiple linear models (mean adjusted-R^2^) = 0.00; all *P*_Imagen_SST_ > 0.05, mean adjusted-R^2^ = -0.02; all *P*_Straitify_MDD_ > 0.05, mean adjusted-R^2^ = 0.00, all *P*_Straitify_AUD_ > 0.05, mean adjusted-R^2^ = 0.01) (see **table S10** for details). The above results hence indicate that CRS is a unique and novel neurocognitive feature.

## Discussion

We demonstrated that integrating brain activation and functional connectivity into an AI-powered Brain-DiT model enables a precise evaluation of personalized neurocognitive processing. Brain-DiT revealed qualitative individual differences in neurocognitive functions, which may represent the neural bases of psychological perceptual or mental sets, *i.e*., subconscious predispositions to perceive information or approach a problem like a task-related fMRI, likely shaped by past experiences and motivations (*27*). Moreover, these neurocognitive profiles had substantial implications for diagnosing relevant mental disorders, such as a perceptual bias for negative stimuli during an emotional task for major depression (*OR* = 9 for high-risk individuals and 11.6 for clinically diagnosed patients) and the lack of strategical adjustment in a motor inhibition task for alcohol use disorders (*OR* = 11.9 for high-risk individuals and 9.2 for clinically diagnosed patients).

A key example of applying Brain-DiT is the identification of the left inferior frontal cortex (left-IFC) as critical for emotional perception in a qualitatively distinct depression subgroup. This finding goes beyond the traditional role of the left-IFC in speech production and semantic retrieval (*37*), as well as in expressing emotional language (*38*). This new understanding may hence imply a shared neurocognitive relationship between emotional and language processing in the left-IFC. Likewise, while alcoholism is often linked to impulsive decision-making (*32*), Brain-DiT modeling of the neurocognitive processes underlying stop-signal task performance suggests an alternative cognitive mechanism (*39*) - a lack of task-related strategy to avoid negative outcomes or aberrant activity in the positive affective system, possibly related to compulsive behavior (*40, 41*). Both findings were supported by causal evidence from lesion maps and brain stimulation.

Thus, the neurocognitive profiles derived from Brain-DiT have revealed findings that were not identified using traditional fMRI measures, such as activation and functional connectivity, alone. This oversight by previous studies may be attributed to two key factors. First, traditional analytical methods in passive paradigms like the emotional face task (EFT) often overlook prior individual differences in perceptual sets, focusing instead on common brain regions implicated in emotional processing (*e.g*., the amygdala). Second, the cognitive relevance score (CRS) derived from Brain-DiT is calculated based on each region’s contribution to a full-brain model, thus capturing the collective effort of brain regions towards performing a task, rather than assessing them in isolation. In summary, contrary to the localized low-dimension traditional approach (*e.g*., the contrast map, **Fig. 1B Left**), Brain-DiT and neurocognitive profiles benefit from the high-dimensional brain-wide modeling of cognitive processing (*i.e*., the distinct transition pathways between brain states according to task conditions; **Fig. 1B Right**).

In conclusion, we propose a new modeling paradigm for personalized neurocognitive profiles in the human brain by applying state-of-the-art diffusion generative models on task-related fMRI data. This approach is able to fully capitalize on large neuroimaging datasets to determine the underlying qualitative neurocognitive biomarkers for a range of mental health disorders, bridging the gap between cognitive neuroscience and psychiatry. By more closely capturing the complexity of individual brain function, the novel Brain-DiT represents a solid step toward truly personalized medicine in psychiatry.

## Acknowledgments

The study was funded by National Key R and D Program of China (2022CSJGG1000 [to T.J.], 2021YFC2501400 [to T.J.], 2019YFA0709501 [to T.J.], 2019YFA0709502 [to J.F.], 2018YFC1312900 [to T.J.] and 2018YFC1312904 [to J.F.]), the National Natural Science Foundation of China (T2122005 [to T.J.], 82150710554 [to G.S.] and 81801773 [to T.J.]), the Shanghai Pujiang Project (18PJ1400900 [to T.J.]), Guangdong Key Research and Development Project (2018B030335001 [to J.F.]), the European Union-funded FP6 Integrated Project IMAGEN (Reinforcementrelated behaviour in normal brain function and psychopathology) (LSHMCT- 2007-037286 [to G.S.]), the 111 Project (B18015 [to J.F.]), the key project of Shanghai Science and Technology (16JC1420402 [to J.F.]), Shanghai Municipal Science and Technology Major Project (2018SHZDZX01 [to J.F.]), Zhangjiang Lab [to J.F.], Shanghai Center for Brain Science and Brain-Inspired Technology [to J.F.]; This work also received support from the following sources: the European Union-funded FP6 Integrated Project IMAGEN (Reinforcement-related behaviour in normal brain function and psychopathology) (LSHM-CT- 2007-037286), the Horizon 2020 funded ERC Advanced Grant ‘STRATIFY’ (Brain network based stratification of reinforcement-related disorders) (695313), Horizon Europe ‘environMENTAL’, grant no: 101057429, UK Research and Innovation (UKRI) Horizon Europe funding guarantee (10041392 and 10038599), Human Brain Project (HBP SGA 2, 785907, and HBP SGA 3, 945539), the Chinese government via the Ministry of Science and Technology (MOST). The German Center for Mental Health (DZPG), the Bundesministerium für Bildung und Forschung (BMBF grants 01GS08152; 01EV0711; Forschungsnetz AERIAL 01EE1406A, 01EE1406B; Forschungsnetz IMAC-Mind 01GL1745B), the Deutsche Forschungsgemeinschaft (DFG project numbers 458317126 [COPE], 186318919 [FOR 1617], 178833530 [SFB 940], 386691645 [NE 1383/14-1], 402170461 [TRR 265], 454245598 [IRTG 2773]), the Medical Research Foundation and Medical Research Council (grants MR/R00465X/1 and MR/S020306/1), the National Institutes of Health (NIH) funded ENIGMA-grants 5U54EB020403-05, 1R56AG058854-01 and U54 EB020403 as well as NIH R01DA049238, the National Institutes of Health, Science Foundation Ireland (16/ERCD/3797). NSFC grant 82150710554. Further support was provided by grants from: - the ANR (ANR-12-SAMA-0004, AAPG2019 - GeBra), the Eranet Neuron (AF12-NEUR0008-01 - WM2NA; and ANR-18-NEUR00002-01 - ADORe), the Fondation de France (00081242), the Fondation pour la Recherche Médicale (DPA20140629802), the Mission Interministérielle de Lutte-contre-les-Drogues-et-les-Conduites-Addictives (MILDECA), the Assistance-Publique-Hôpitaux-de-Paris and INSERM (interface grant), Paris Sud University IDEX 2012, the Fondation de l’Avenir (grant AP-RM-17-013), the Fédération pour la Recherche sur le Cerveau; ImagenPathways “Understanding the Interplay between Cultural, Biological and Subjective Factors in Drug Use Pathways” is a collaborative project supported by the European Research Area Network on Illicit Drugs (ERANID). This paper is based on independent research commissioned and funded in England by the National Institute for Health Research (NIHR) Policy Research Programme (project ref. PR-ST-0416-10001). The views expressed in this article are those of the authors and not necessarily those of the national funding agencies or ERANID. This work also received support from the following sources: the Horizon 2020 funded ERC Advanced Grant ‘STRATIFY’ (Brain network based stratification of reinforcement-related disorders; 695313), the Medical Research Council and Medical Research Foundation (grants MR/R00465X/1 and MRF-058-0004-RG-DESRI: ‘ESTRA: Neurobiological underpinning of eating disorders: integrative biopsychosocial longitudinal analyses in adolescents’; MR/S020306/1 and MRF-058-0009-RG-DESR-C0759: ‘Establishing causal relationships between biopsychosocial predictors and correlates of eating disorders and their mediation by neural pathways’) and the National Institute for Health and Research (NIHR) Biomedical Research Centre (BRC) and Maudsley NHS Foundation Trust (SLaM).

## Author Contributions

Tianye Jia, Weikang Gong, and Jianfeng Feng conceptualized the study. Rongquan Zhai, Liping Zheng, Weikang Gong, and Tianye Jia designed the analytic approach; Danilo Bzdok helped to review and ensure the validity of the machine learning approaches; Rongquan Zhai, Liping Zheng, and Yechen Hu analyzed the data; Rongquan Zhai and Shitong Xiang helped in organizing the original behavioral scale and preprocessing the imaging data; Rongquan Zhai and Tianye Jia drafted the manuscript; Weikang Gong, Edmund T. Rolls and Trevor W. Robbins revised the first draft; Tianye Jia, Weikang Gong, Chao Xie, Edmund T. Rolls, Barbara Sahakian and Trevor W. Robbins interpreted the results; Rongquan Zhai, Liping Zheng, Lei Peng and Yechen Hu contributed to figure visualization; Tobias Banaschewski, Gareth J. Barker, Arun L.W. Bokde, Rüdiger Brühl, Sylvane Desrivières, Herta Flor, Hugh Garavan, Penny Gowland, Antoine Grigis, Andreas Heinz, Herve Lemaitre, Jean-Luc Martinot, Marie-Laure Paillère Martinot, Eric Artiges, Frauke Nees, Dimitri Papadopoulos Orfanos, Tomáš Paus, Luise Poustka, Michael N. Smolka, Sarah Hohmann, Nathalie Holz, Nilakshi Vaidya, Robert Whelan, Gunter Schumann, Henrik Walter and Tianye Jia were the principal investigators of IMAGEN Consortium and helped in data acquirement. Nilakshi Vaidya, Zuo Zhang, Lauren Robinson, Jeanne Winterer, Sinead King, Yuning Zhang, Gareth J Barker, Arun L Bokde, Rüdiger Brühl, Hedi Kebir, Hervé Lemaître, Frauke Nees, Dimitri Papadopoulos, Ulrike Schmidt, Julia Sinclair, Argyris Stringaris, Robert Whelan, Henrik Walte, Sylvane Desrivières, Gunter Schumann were the principal investigators of STRATIFY Consortium and helped in data acquirement.

## IMAGEN Consortium Authors

Tobias Banaschewski, Gareth J. Barker, Arun L.W. Bokde, Rüdiger Brühl, Sylvane Desrivières, Herta Flor, Hugh Garavan, Penny Gowland, Antoine Grigis, Andreas Heinz, Herve Lemaitre, Jean-Luc Martinot, Marie-Laure Paillère Martinot, Eric Artiges, Frauke Nees, Dimitri Papadopoulos Orfanos, Tomáš Paus, Luise Poustka, Michael N. Smolka, Sarah Hohmann, Nathalie Holz, Nilakshi Vaidya, Henrik Walter, Robert Whelan, Gunter Schumann and Tianye Jia.

## STRATIFY Consortium Authors

Nilakshi Vaidya, Zuo Zhang, Lauren Robinson, Jeanne Winterer, Sinead King, Yuning Zhang, Gareth J Barker, Arun L Bokde, Rüdiger Brühl, Hedi Kebir, Hervé Lemaître, Frauke Nees, Dimitri Papadopoulos, Ulrike Schmidt, Julia Sinclair, Argyris Stringaris, Robert Whelan, Henrik Walte, Sylvane Desrivières, Gunter Schumann.

## Data Availability

IMAGEN data are available from a dedicated database at https://imagen2.cea.fr. Written and informed consent was obtained from all participants by the IMAGEN consortium and the study was approved by the institutional ethics committee of King’s College London (PNM/10/11-126), University of Nottingham (D/11/2007), Trinity College Dublin (SPREC092007-01), Technische Universitat Dresden (EK 235092007), Commissariat a l’Energie Atomique et aux Energies Alternatives, INSERM (2007-A00778-45), University Medical Center at the University of Hamburg (M-191/07) and in Germany at medical ethics committee of the University of Heidelberg (2007-024N-MA) in accordance with the Declaration of Helsinki. The details of the Stratify project could be found at https://stratify-project.org/. The imaging protocols were harmonized to match with the IMAGEN protocols. The task-based MRI data were also collected in the Stratify project. The paradigms of the emotional faces task and stop-signal task were the same as those in the IMAGEN project. The data processing procedures were the same as those for the IMAGEN project. The IMAGEN project data are available from a dedicated database: https://imagen-project.org. The Stratify project data are available from a dedicated database: https://stratify-project.org/.

## Code Availability

The code that supports the findings of this study will be made available in the future version.

## Supplementary Materials

Materials and Methods

Supplementary Text

Figs. S1 to S6

Table S1 to S11

