## Supplementary Materials for "Brain Diffusion Transformer for Personalized Neuroscience and Psychiatry"

**The PDF file includes:**

Materials and Methods
Supplementary Text
Figs. S1 to S7
Table S1 to S11
References

### Materials and Methods

In this section, we summarize overviews of datasets, experimental designs for fMRI tasks, protocols for multimodal brain imaging, behavioral questionnaires, and the diffusion generative model implemented in the present study. Furthermore, we detailed the computational processes of establishing the personalized neurocognitive relevance score map based on HCPex(*1*) template through digital lesioning.

#### Experiment designs of the IMAGEN cohort and Multiple Modalities Data preprocessing

##### *Exploratory Dataset*

IMAGEN (*2, 3*), a comprehensive longitudinal neuroimaging-genetics cohort study, aims to uncover the biological foundations of individual differences in psychological and behavioral traits and their links to common psychiatric disorders. Initially enrolling 2,000 participants (a population cohort) at age 14, with 1,300 retained at age 19, the study conducts extensive neuropsychological, behavioral, clinical, and environmental assessments. It also gathers T1-weighted and diffusion-weighted structural MRI and task-based and resting-state fMRI. The present investigation specifically utilized task and resting-state fMRI as well as behavioral data. As a population-based approach, IMAGEN notably maintains balanced sample sizes between male and female participants, based on self-reported sex.

##### *Validation Clinic Dataset*

The Stratify project recruited patients (ages 19–25) with alcohol use disorder (AUD) and major depressive disorder (MDD), along with controls at three sites: Berlin, London, and Southampton (*4*). The Stratify protocol was harmonized to match the IMAGEN protocol. Similarly, the Stratify datasets also collected task-based fMRI data for the EFT and SST, as well as resting fMRI. After quality control to match for sites and sex between cases and controls, there are 127 MDD patients and 66 controls for the EFT, as well as 119 AUD patients and 61 controls for the SST. The fMRI experimental paradigms used in STRATIFY are consistent with those employed in the IMAGEN study.

##### *Emotional Face Task*

IMAGEN conducted a modified emotional face task (EFT) (*5*). The task presents participants with alternating 18-second blocks of emotive facial displays and control stimuli. The facial stimuli comprised brief (200-500ms) monochromatic video clips of male and female faces expressing anger, neutrality, or happiness. The experiment included four blocks for each emotion, interspersed with 12 control blocks, each emotion block containing eight trials featuring six distinct facial identities with equal gender representation. Control blocks displayed monochromatic concentric circles expanding and contracting to match the contrast and motion characteristics of the facial stimuli. Our analysis concentrated on responses to angry, neutral, and happy expressions, chosen for their widespread recognition and frequent use in emotional response studies. The task's structure and stimulus selection, grounded in established research methodologies, aimed to effectively evoke and measure emotional responses, providing a controlled environment for investigating emotional processing in response to socioemotional visual stimulation. This approach contributes to the broader understanding of emotional cognition while building upon previous work in the field, ensuring reliability and reproducibility. The same EFT was also conducted in the Stratify cohort.

##### *Stop-signal Task*

IMAGEN also conducted an event-related Stop-signal task (SST) (*6*) to examine neural correlates of inhibitory control, building on established cognitive neuroscience methodologies. The experimental paradigm comprised go trials (83%, n=480) and stop trials (17%, n=80). During go trials, participants responded to directional arrows with corresponding finger presses, while stop trials introduced an additional upward arrow, signaling response inhibition. A dynamic tracking algorithm modulated the stop-signal delay based on cumulative performance, aiming for a theoretical 50% inhibition success rate and challenging participants at their inhibitory threshold. With a fixed 1,800 ms inter-trial interval, the task design elicited balanced successful and unsuccessful inhibition trials. Our analysis focused on two key measures: stop success and go success, enabling a comprehensive examination of inhibitory processes and overall task performance. This experimental framework, grounded in cognitive control theory, provides a robust approach for investigating the neural mechanisms underlying response inhibition and cognitive flexibility in dynamic task environments, contributing to our understanding of these crucial cognitive processes, which is complementary to EFT in experimental design (event-related vs. block design; proactive response vs. passive viewing). The same SST was also adopted in the Stratify cohort.

##### *Image Acquisition*

Neuroimaging data for this study were acquired at eight IMAGEN assessment centers using 3-Tesla MRI systems from diverse manufacturers (Siemens, Philips, General Electric, and Bruker). To ensure cross-site compatibility, we implemented a uniform scanning protocol with parameters optimized for all scanner types. The imaging sequence included high-resolution T1-weighted 3D structural scans for anatomical detail and co-registration, alongside functional images obtained using gradient-echo, echo-planar imaging (EPI) to measure blood oxygen level-dependent (BOLD) signals. For all functional tasks, we used a consistent configuration of 40 slices aligned to the anterior commissure-posterior commissure (AC-PC) line, with 2.4 mm slice thickness and 1 mm gap. Signal quality was optimized, particularly for subcortical regions, using an echo time of 30 ms and a repetition time of 2,200 ms. This carefully designed multi-site protocol facilitated the collection of high-quality structural and functional data, ensuring consistency and comparability across all centers - a crucial factor for robust analyses in large-scale neuroimaging studies.

#### *Task-based Functional Image Preprocessing*

Preprocessing of task-based functional MRI data was conducted using Statistical Parametric Mapping software (SPM8). The preprocessing pipeline encompassed several steps to enhance data quality and prepare it for subsequent analyses. Initially, slice timing correction was applied to account for temporal discrepancies arising from multi-slice acquisition. This was followed by realignment of all volumes to the first acquired image to mitigate motion-related artifacts. Spatial normalization was then performed, warping the data nonlinearly to a custom echo-planar imaging template in Montreal Neurological Institute (MNI) space. This template, with dimensions of 53 × 63 × 46 voxels, was derived from the mean images of 400 adolescent participants, ensuring age-appropriate registration. The normalized data were resampled to a 3 × 3 × 3 $mm^3$ voxel resolution. Finally, to improve the signal-to-noise ratio and accommodate inter-subject variability, the brain images were spatially smoothed using an isotropic Gaussian kernel with a 5 mm full-width at half-maximum(*7*). This comprehensive preprocessing approach aimed to optimize the data for robust statistical analyses while maintaining spatial specificity.

#### *Patch-based Resting-state Functional Connectivity*

Resting-state functional connectivity (rsFC) in this study was computed using a patch-based parcellation approach to train the diffusion transformer model. First, the original 3D spatial dimensions of the resting-state fMRI data were resliced into a 64 × 64 × 64 grid of voxels, preserving the temporal dimension. This grid was then regrouped into 512 non-overlapping 8 × 8 × 8 patches (i.e., merging every 8 voxels into 1 patch along each spatial axis) to reduce computational burdens. For each patch, we calculated a temporal correlation matrix using Pearson’s correlation coefficient between this patch’s time series and the time series of each of the 512 patches across the entire brain for each participant. This produced 512 unique functional connectivity (FC) values per patch, organized as an 8 × 8 × 8 matrix representing the patch’s connectivity profile Fig S7.

By repeating this process across all 512 patches (in total 512 × 512 = 64 × 64 × 64 entries), we constructed a hierarchical hypermatrix with two nested layers:

Outer layer: An 8 × 8 × 8 matrix where each entry corresponds to the spatial location of a patch.

Inner layer: For each outer-layer entry, an 8 × 8 × 8 matrix encoding the FC values that quantify the patch's whole-brain communication patterns.

When integrated with the iterative generative processes of diffusion transformers—which progressively refine noise into structured outputs—this nested hypermatrix structure may approximate aspects of recursive information processing observed in the brain, such as hierarchical feedback loops, multi-scale integration, and dynamic reconfiguration of functional networks (*8*). The detailed calculation framework can be seen in Fig. S7.

#### *Voxel-based Morphometry*

Structural T1-weighted images were processed using the Statistical Parametric Mapping (SPM) software package, developed by the Wellcome Department of Neuroimaging in London, UK. This analysis was conducted within the MATLAB environment (version R2020b). To perform voxel-based morphometry (VBM), we employed the VBM8 toolbox, an extension of SPM. The initial step involved segmentation of the images into three tissue classes: gray matter, white matter, and cerebrospinal fluid. Subsequently, we utilized the gray matter segments to construct a study-specific template through the DARTEL (Diffeomorphic Anatomical Registration Through Exponentiated Lie Algebra(*9*)) algorithm. The gray matter maps were then spatially normalized to the Montreal Neurological Institute (MNI) standard space, with a resolution of 1.5 × 1.5 × 1.5 mm. This normalization process incorporated both linear and non-linear transformations, and the resulting maps were modulated by applying the Jacobian determinants derived from these transformations. In the final preprocessing step, the normalized and modulated gray matter maps underwent spatial smoothing using an 8 mm full-width at half-maximum (FWHM) Gaussian kernel. The resulting voxel values in these processed maps represent local gray matter volume (GMV) proportions.

#### *Patch-based Diffusion Tensor Image*

Diffusion-weighted imaging (DWI) data were preprocessed using a combination of FSL version 6.0.4 (FMRIB Software Library) and MRtrix3 (version 3.0) software packages. The preprocessing pipeline included several crucial steps to enhance data quality and correct for various artifacts, including denoising, Gibbs ringing artifact correction, and comprehensive corrections for head motion, eddy current-induced distortions, and susceptibility-induced off-resonance geometric distortions utilizing reversed phase-encoding b=0 s/mm$^2$ images. Additionally, the bias field arising from nonuniform coil receive sensitivity was mitigated. To construct the anatomical connectivity matrix, we first generated a seed mask delineating the gray matter-white matter interface. Whole-brain white matter tractography was then performed, employing this seed mask to initiate streamline propagation. The resulting tractogram was used to quantify the number of fiber tracts connecting each pair of voxels within the mask, thereby populating the anatomical connectivity matrix.

Diffusion Tensor Imaging (DTI) in this study was computed using a patch-based hypermatrix framework analogous to our rs-FC approach. First, the imaging space was parcellated into an 8 × 8 × 8 grid, creating 512 non-overlapping cubic patches. This generated a novel patch-wise structural template optimized for computational efficiency and spatial coherence.

For each of the 512 × 512 patch pairs, we calculated structural connectivity matrices using DTI-derived parameters (e.g., fractional anisotropy, axial/radial diffusivity) if both patches contained brain tissues (239 × 239 pairs), while the rest of the patch pairs were set zero. These matrices were then embedded into a hierarchical hypermatrix structure with two nested layers:

Outer layer: An 8x8x8 matrix encoding the spatial coordinates of each patch.

Inner layer: For each outer-layer entry, an 8 × 8 × 8 matrix storing structural connectivity values between the target patch and all brain patches.

This approach enables direct comparison with the aforementioned rs-FC hypermatrices, given the identical form of the final data representation.

#### *DAWBA and SDQ*

The behavioral symptoms of IMAGEN participants were assessed using screening questions from the Development and Well-Being Assessment (DAWBA) (*12*) and the Strengths and Difficulties Questionnaire (SDQ) (*13*) at three data collection timepoints (ages 14, 19, and 23). DAWBA is a comprehensive psychiatric screening questionnaire. It has previously been used with success to define subthreshold clinical symptoms in neuroimaging studies of subclinical psychopathology (*14*). The SDQ was also employed in the current investigation, as it aids in assigning diagnostic status within the DAWBA framework. Specifically, we included self-rated (*15*) emotional problems from the SDQ in this study, and also combined 8 items from SDQ and DAWBA to form the depression symptoms (*16*). Additionally, DAWBA provides a diagnostic output compatible with ICD10/DSM5 for common psychiatric disorders based on the likelihood of a clinical diagnosis following the ratings. In this study, we considered individuals with ratings greater than 3 as high-risk (i.e., having over a 50% chance of being diagnosed).

#### *TCI-R and SURPS*

In the IMAGEN cohort, impulsivity as a personality trait was measured using the Temperament and Character Inventory-Revised (TCI, 36 items) (*17*) and the Substance Use Risk Personality Scale (SURPS, 23 items, self-questionnaire) (*18*) across all three timepoints (ages 14, 19, and 23).

##### *AUDIT*

The alcohol abuse was assessed using the screening questions from the Alcohol Use Disorders Identification Test (AUDIT, 10 items)(*19*). The AUDIT was developed by the World Health Organization as a simple way to screen and identify people who are at risk of developing alcohol problems. The AUDIT test focuses on identifying the preliminary signs of hazardous drinking and mild dependence. It is used to detect alcohol problems experienced within the last year. In the present study, we used the total AUDIT score as a measure of alcohol abuse behavior in the population and "total score >= 8" as an indicator of high risk for alcohol dependence.

##### *BIS*

The Barratt Impulsiveness Scale-11 (BIS-11)(*20*) is a 30-item self-report questionnaire that assesses three dimensions of impulsivity: attentional, motor, and non-planning. It uses a 4-point Likert scale and is widely employed in clinical and research settings to evaluate impulsive traits.

##### *CGT*

The Cambridge Gambling Task (CGT) is a neuropsychological assessment tool within the Cambridge Neuropsychological Test Automated Battery (CANTAB) designed to evaluate risk-based decision-making and impulsivity. It quantifies how individuals balance potential rewards against risks and assesses their ability to adjust strategies under varying conditions. The CGT quality of decision-making is the proportion of trials on which the subject chooses the most likely outcome. The CGT deliberation time is the reaction time to choose the color of the box. The overall bet is the overall bet across the trials. The CGT risk-taking: mean proportion of available points the subject stakes at each trial. The CGT delay aversion is the difference between the risk-taking score in the descending and the ascending conditions. The CGT risk adjustment is the degree to which a subject adjusts the risk taking according to the ratio of colored boxes, calculated as: [2 x (proportion of pints staked (%) at 9:1) + (% 8:2) - (% 7:3) – 2 x (% 6:4)] / CGT risk-taking.

##### *Affective Go/No-Go Task*

The Affective Go/No-go (AGN) task (*21*) is part of the Cambridge Cognition Battery (CANTAB, http://www.cambridgecognition.com/). The AGN assesses the interplay between emotional processing and executive function, specifically evaluating how emotional stimuli influence inhibitory control (the ability to suppress prepotent responses). It is widely used to study cognitive-emotional dysregulation in psychiatric disorders (e.g., depression, anxiety, bipolar disorder). The calculated measures are mean correct latency for positive and negative stimuli, and number of omission errors for positive and negative stimuli. We also investigated for emotional bias between positive and negative stimuli for both response latency and omission errors.

#### Diffusion Generative Model

As a class of probabilistic models used for generating data by reversing a diffusion process (*22*), a diffusion generative model starts with a simple distribution and progressively transforms it into a complex target distribution through a series of steps. The input for these models typically consists of a noisy version of the data or Gaussian noise, which is refined progressively through a denoising process. The output is the generated data that closely resembles the target distribution. In the present study, the input includes brain activations during a first task condition along with additional information such as rsFC or structural MRI measures like VBM and DTI, and the output is the brain activation during the target second task condition, and a hence established generative model tracks brain's neurocognitive state transitions between task conditions, constrained by an additional non-task brain feature, such as rsFC, VBM or DTI. Particularly, we employed the U-ViT (U-shaped Vision Transformer) (*23*) model, which is based on the Transformer (*24*) with the blocks modified to fit a 3D denoising generative architecture.

##### *Construction of noise-to-activation map generator—model training phase*

The U-ViT model is trained using a noise prediction objective function. It takes time $t$, conditional information $c$, and noisy image $x_t$ as inputs, treating them all as tokens fed into the Transformer structure. The model learns by minimizing the mean squared error between the predicted noise $\boldsymbol{\epsilon_\theta}$ and the actual noise $\boldsymbol{\epsilon}$ added. Key design elements include long skip connections between shallow and deep layers, as well as an optional 3x3x3 convolutional block before the output.

In detail, we aimed to minimize the following loss function:

$$\min_{\boldsymbol{\theta}} \mathbb{E}_{t,\boldsymbol{x}_0,c,\boldsymbol{\epsilon}} \|\boldsymbol{\epsilon} - \boldsymbol{\epsilon_\theta}(\boldsymbol{x}_t, t, c)\|_2^2,$$

By minimizing this loss function, we could accurately predict the noise added at each step, thereby achieving denoising and ultimately accomplishing the generation objective.

##### *Personalized activation map generation from noise—model inference phase*

During inference, the U-ViT model starts from pure noise and gradually denoises to generate an image. At each step, the inputs include the current timestep $t$, conditional information (*e.g.,* task activation and other functional or structural measurements in the present study), and the current noisy image. The model will predict the noise, which is then used to update the current noisy image. This process iterates backward from high to low noise, eventually producing a clear image sample. The entire process utilizes the learned noise prediction capability to progressively refine and improve image quality. During the inference stage, we uniformly set the number of inference steps to 50, as the generation performance tends to stabilize thereafter.

##### *Implementation Details*

We stratified the IMAGEN cohort participants according to gender and site information (with two genders and eight sites, resulting in 16 subgroups), ensuring a balanced data distribution. We selected 70% of each class as the training set, 10% as the validation set, and the remaining 20% as the hold-out test set. Due to the longitudinal nature of the data, we also ensured that all individuals had data presented only in either the training, validation, or testing set.

The model was trained on one A100 card for 6.5 hours, and the inference was fast, *i.e.,* 10 minutes for 200 subjects. Our model parameters are 38 million in total. Detailed model parameters can be found in **table S11**. In this study, participants were split once into training (70%), validation (10%), and testing (20%) sets, with careful stratification by gender, research site, and longitudinal timepoints to ensure balanced representation. The Brain-DiT model was trained and fine-tuned on the training and validation sets, and its performance was evaluated on the held-out test set. The reported performance metrics, such as Pearson's r, were calculated based on this single split.

#### Comparison of lesioning effects between different masks

During the process of systematically lesioning the brain (**Fig. 3 B&H**), a specific algorithm for comparing three different masks (*i.e.,* the individual activation map (I-map), the Neurosynth meta map (N-map, starting from 50%), and the group-level mean activation map (G-map)) was adopted: for each individual, the prediction accuracy across all proportions was summed up, and paired t-tests were applied to compare the summed prediction accuracy between the three masks. Notably, when the N-map was used in the comparison, the proportions included were 50%, 30%, and 10%, instead of the full set (*i.e.,* 90%, 70%, 50%, 30%, or 10%).

#### Computation of individual cognitive relevance map

Each individual receives a cognitive relevance map for the EFT or SST task, which evaluates the contribution of each brain region to the transformation performance (or prediction accuracy). Specifically, the calculation starts with a trained diffusion generative model from the training set. During inference for each individual in the test set, based on a given template (e.g., HCPex (*1, 25*) in the present study), each brain region will be removed from the activation map, and the corresponding decrease (compared to the use of whole brain activation map) in the spatial similarity between the predicted and observed brain activation map will be recorded as the cognitive relevance score (CRS):

$$crs_i = r_{WB} - r_{WB/i},$$

where $crs_i$ is the cognitive relevance score of the selected brain region $i$, $r_{WB}$ is the prediction similarity using the whole brain activation map, and the $r_{WB/i}$ is the prediction similarity using the data except for the brain region i. A participant's cognitive relevance map is then obtained by exploring the entire brain.

#### Individual clustering by cognitive relevance map

We first organized all participant's cognitive relevance maps into a two-dimensional matrix, where the rows represent subjects, and the columns represent the cognitive relevance scores across all brain regions. Then, we used the R package NbClust (*26*) to perform the hierarchical clustering, where the optimal number of clusters and the clustering outcome will be automatically calculated.

#### Detailed index of key brain regions in HCPex(*1*) atlas

LIFC: 161,162,163,164,165,168,169,170,171,172

RdACC: 323,324,325,321,322

RSTS: 243,242,245,246,241

L/RV1:1,2,3,4,5,6,7,8,9,10,180,181,182,183,184,185,186,187,188,189,190

pgACC: 137,138,139,140,141,324,323,321,325

SMA: 40,39,38,220,221,219

mPFC: 146,147,148,149,150,151,152

L/RV1: 1,3,5,7,180,182,185,187

### Supplementary Text

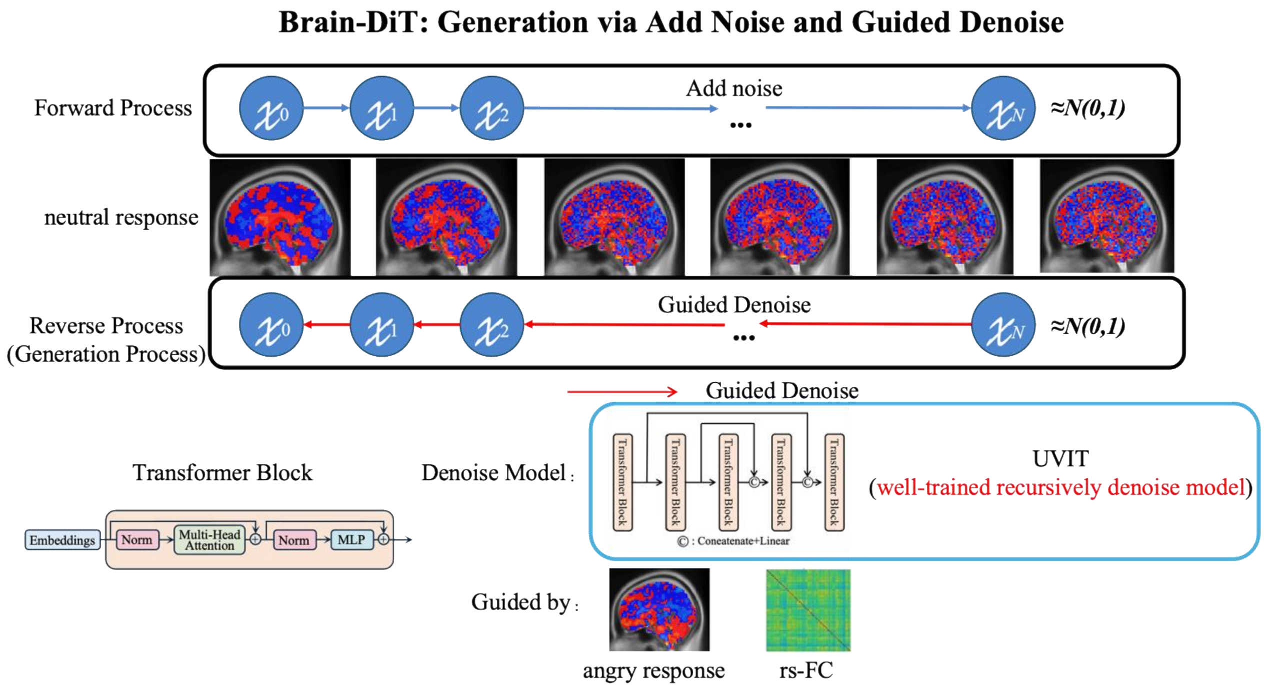


**Fig. S1 The Brain Diffusion Transformer (Brain-DiT) achieves the generation ability of brain activation images through recursive denoising.**

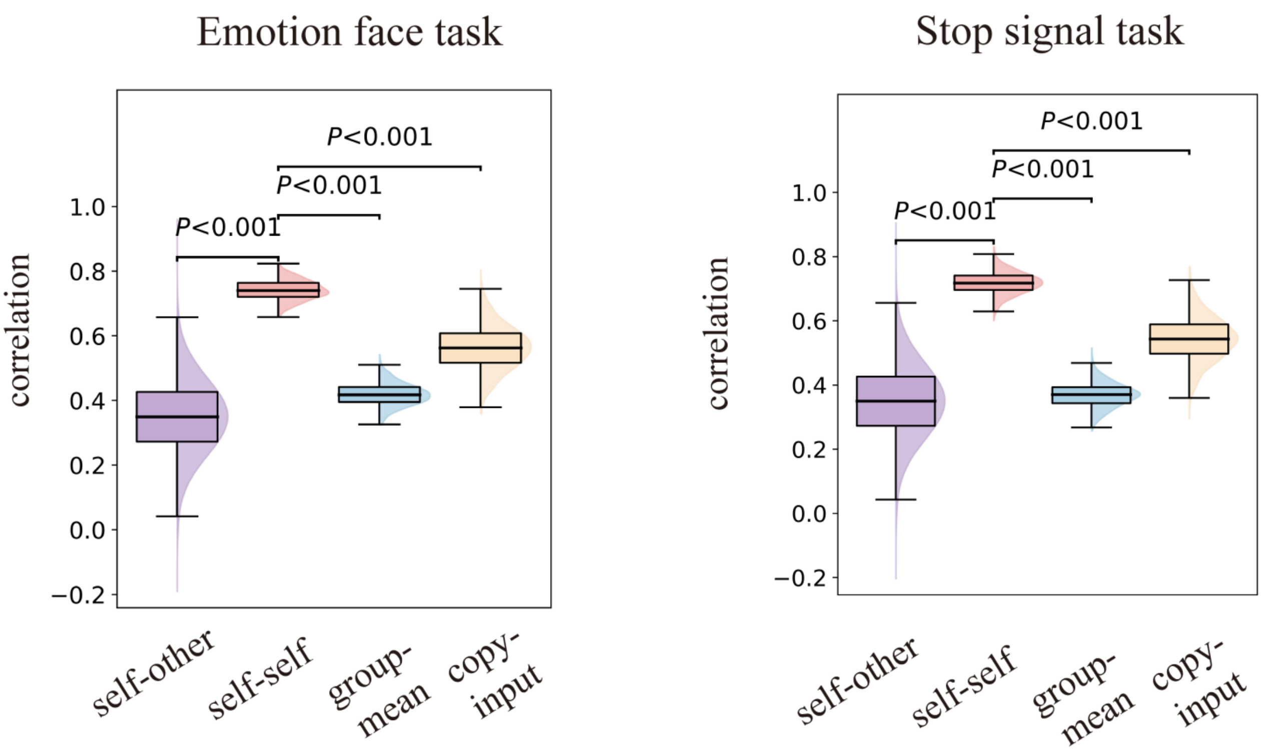


**Fig. S2 Individualized Prediction Ability of Brain-DiT.**

In both emotional face task and stop-signal task, within-subject prediction-actual correlations (labeled self-self; the diagonal elements of the matrix in Fig. 2 lower graphs for A and C) are significantly higher than cross-subject prediction-actual correlations (labeled self-other; the off-diagonal elements of the matrix in Fig. 2 lower graphs for A and C), as well as higher than those by taking the group mean activation (labeled group-mean) or the input signal (labeled copy-input) as the prediction.

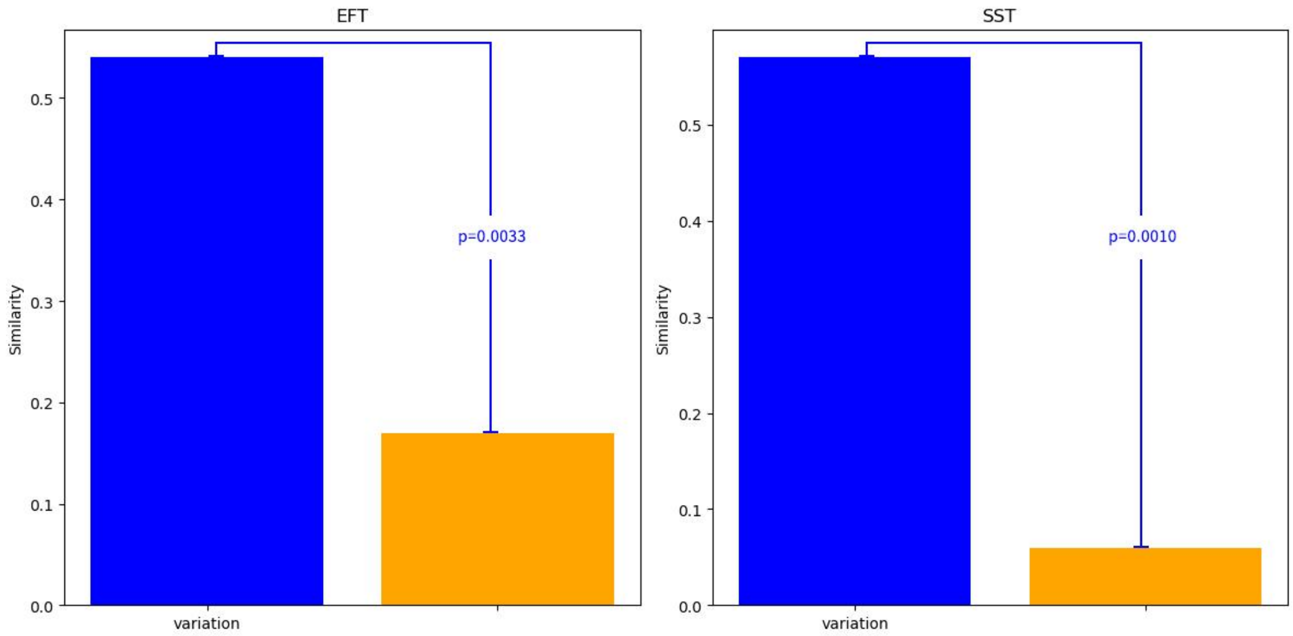


**Fig. S3 The voxel-wise prediction-actual similarity map has a higher spatial similarity with the voxel-wise variation map than with the activation T-map (*i.e.*, the group mean activation).**

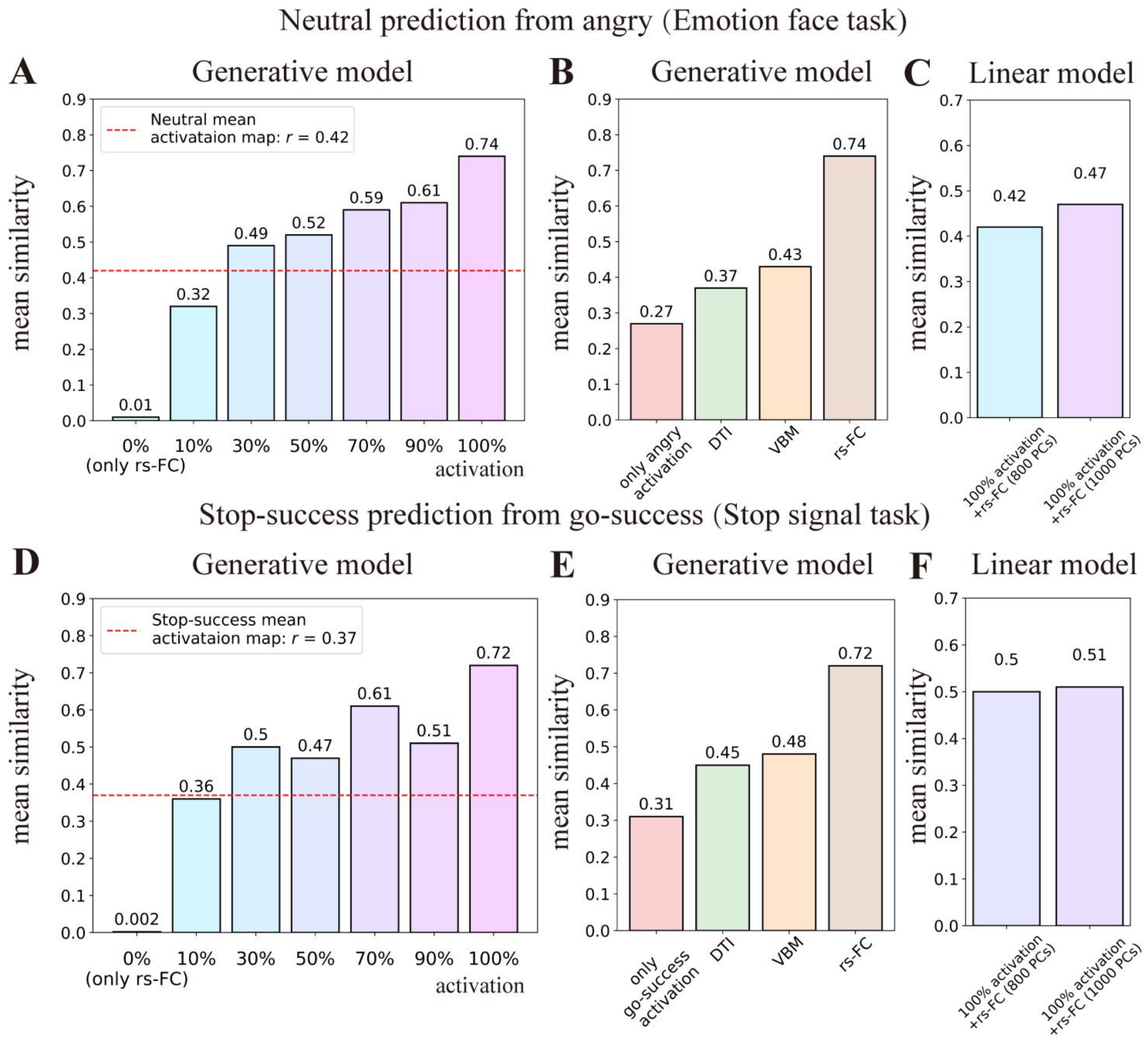


**Fig. S4 Effects of activation map proportions and imaging modalities on the hence-trained generative models' performance and the comparison with linear models' performance. A.** The prediction performance (spatial correlations with the neutral face mean activation in the testing sample) of generative cognitive diffusion transformers (Brain-DiTs) trained with different proportions of the angry face activation map. **B.** The prediction performance (spatial correlations with the neutral face activation in the testing sample) of generative models trained with different MRI modalities (in addition to the activation map of the angry face condition). **C**. The prediction performance for the neutral face activation with different settings of the linear model SuperbigFLICA(*27*). **D.** The prediction performance (spatial correlations with the go-success mean activation in the testing sample) of generative cognitive diffusion transformers (Brain-DiTs) trained with different proportions of the stop-success activation map. **E.** The prediction performance (spatial correlations with the go-success activation in the testing sample) of generative models trained with different MRI modalities (in addition to the activation map of the stop-success condition). **F.** The prediction performance for the go-success activation with different settings of the linear model SuperbigFLICA(*27*).

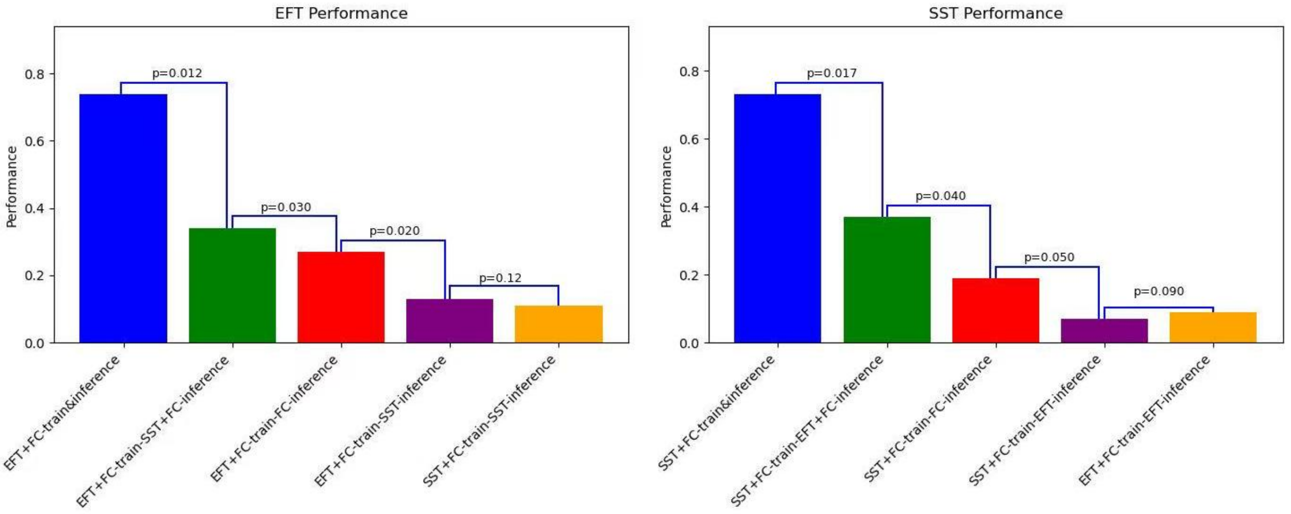


**Fig. S5 Generation performance based on different combinations of fMRI modalities for EFT (Left) and SST (Right).**

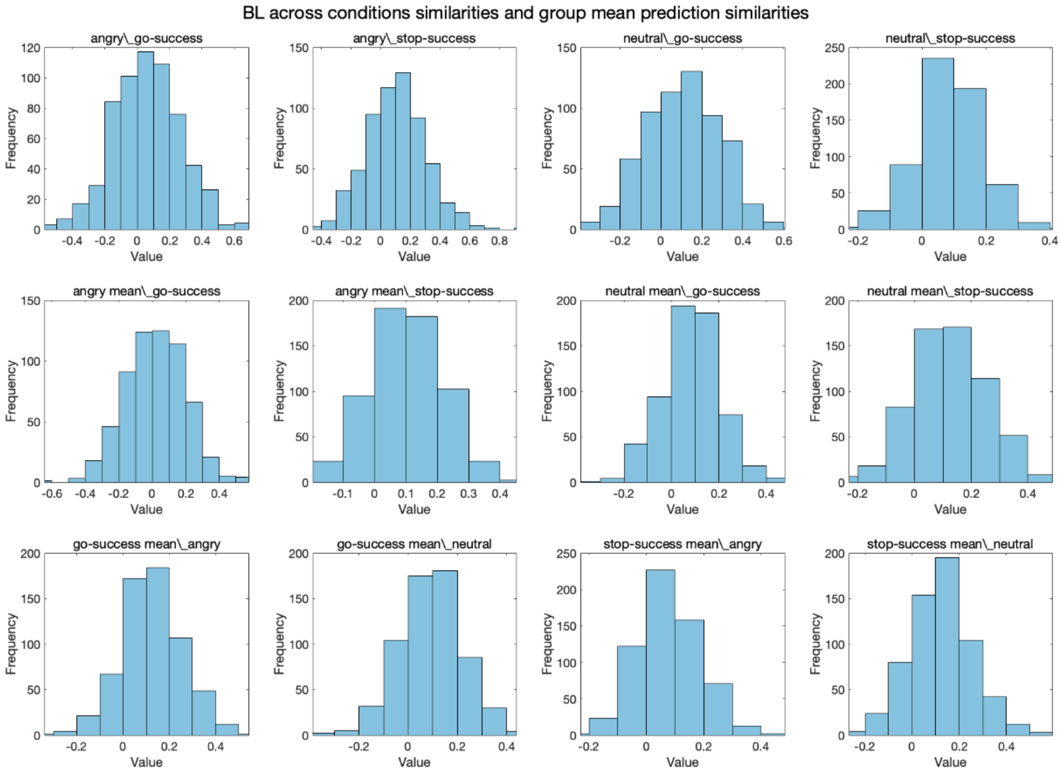
BL across conditions similarities and group mean prediction similarities
angry_go-success
angry_stop-success
neutral_go-success
neutral_stop-success
angry mean_go-success
angry mean_stop-success
neutral mean_go-success
neutral mean_stop-success
go-success mean_angry
go-success mean_neutral
stop-success mean_angry
stop-success mean_neutral
Frequency
Value

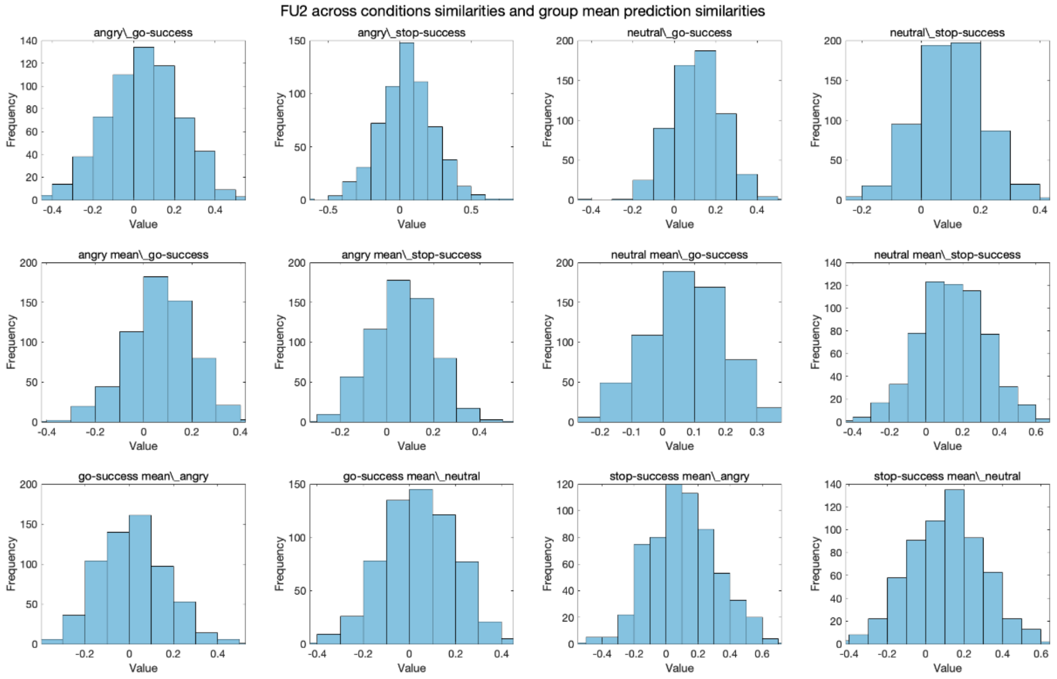
FU2 across conditions similarities and group mean prediction similarities
angry_go-success
angry_stop-success
neutral_go-success
neutral_stop-success
angry mean_go-success
angry mean_stop-success
neutral mean_go-success
neutral mean_stop-success
go-success mean_angry
go-success mean_neutral
stop-success mean_angry
stop-success mean_neutral
Frequency
Value

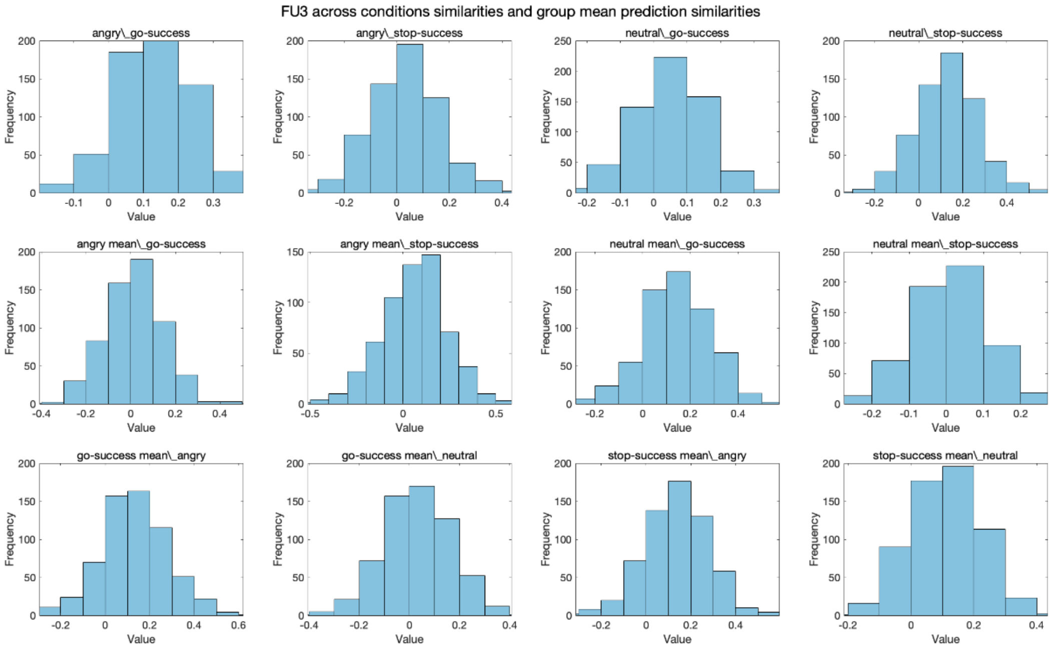


**Fig. S6 Distributions of cross-task similarities between EFT and SST in BL, FU2, and FU3.**

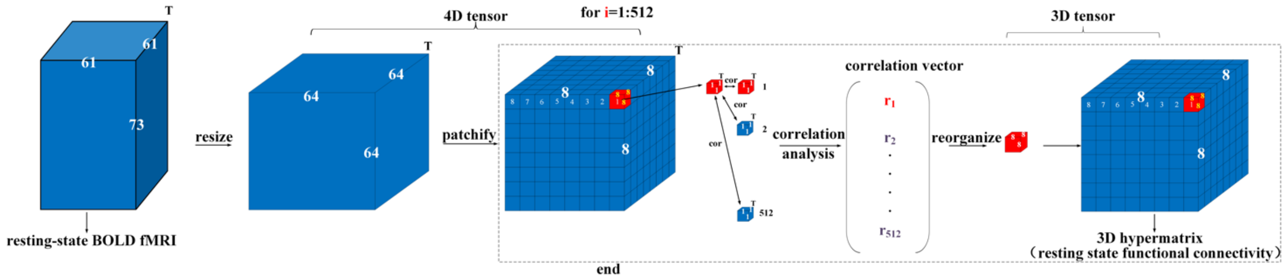


**Fig. S7 The calculation framework of patch-based resting-state functional connectivity.**

**Table S1. Silhouette coefficients and optimal cluster determination in hierarchical clustering based on cognitive relevance scores (CRS).**

(A) Dendrograms for the emotion face task (EFT; upper row) and stop-signal task (SST; lower row) at baseline (BL; first column), and 2-year (FU2; second column) and 3-year (FU3; third column) follow-ups. (B) Silhouette coefficients suggest two-cluster solutions optimal (highlighted in red) for both EFT and SST across all timepoints (the highest coefficient for each model was highlighted in bold).

(A)

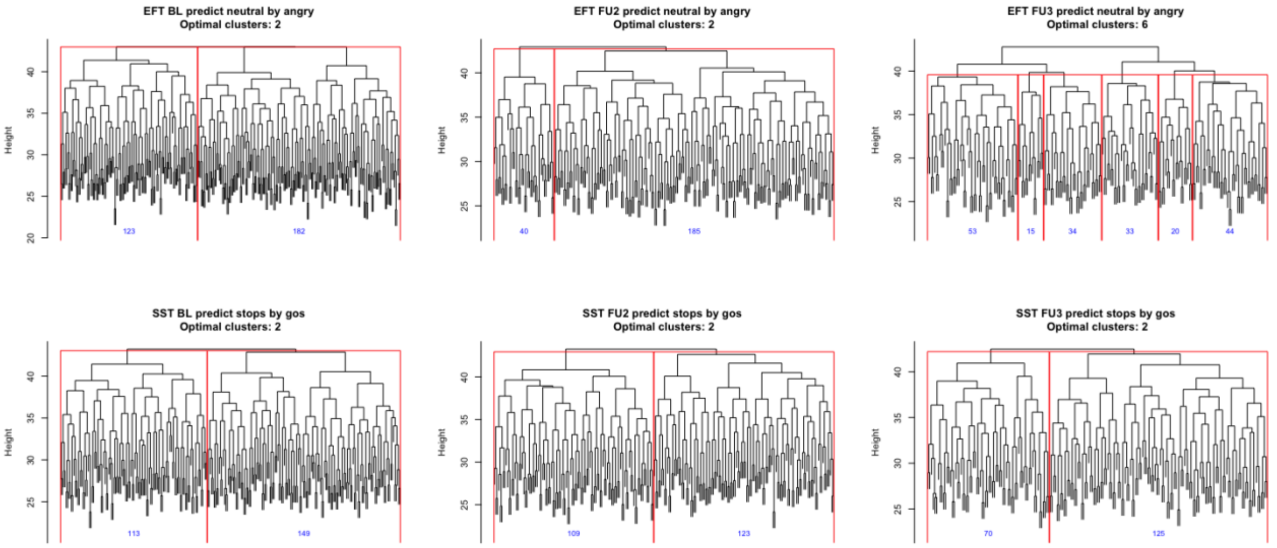


(B)

| BL | **2** | 3 | 4 | 5 | 6 |
|---|---|---|---|---|---|
| EFT | **0.52** | 0.48 | 0.35 | 0.3 | 0.42 |
| SST | **0.6** | 0.35 | 0.39 | 0.46 | 0.51 |

| FU2 | **2** | 3 | 4 | 5 | 6 |
|---|---|---|---|---|---|
| EFT | **0.56** | 0.43 | 0.22 | 0.17 | 0.27 |
| SST | **0.63** | 0.46 | 0.30 | 0.20 | 0.11 |

| FU3 | **2** | 3 | 4 | 5 | 6 |
|---|---|---|---|---|---|
| EFT | 0.56 | 0.49 | 0.31 | 0.39 | **0.61** |
| SST | **0.66** | 0.39 | 0.41 | 0.46 | 0.50 |

**Table S2. Longitudinal stability of neurocognitive subgroups in IMAGEN cohort.**

(A) Subgroups of EFT based on hierarchical clustering results from Table S1. EFT subgroups (left-IFC-dominant vs. dACC/V1-dominant) have a low BL-FU2 stability (13%) but high FU2-FU3 stability (75%). (B) Subgroups of SST based on hierarchical clustering results from Table S1. SST subgroups (high vs. low engagement of pgACC/mPFC/SMA) maintained a high BL-FU3 stability (BL-FU2 80% and FU2-FU3 58%). Abbreviations: EFT, emotional face task; SST, stop-signal task; IFC, inferior frontal cortex; dACC, dorsal anterior cingulate cortex; V1: primary visual cortex; pgACC, pregenual anterior cingulate cortex; mPFC, medial prefrontal cortex; SMA, supplementary motor area; BL, baseline; FU2, follow-up 2; FU3, follow-up3.

(A)

**EFT clustering results**

| | BL | FU2 | FU3 |
|---|---|---|---|
| Group A | 123 | 40 | 102 |
| Group B | 182 | 185 | 97 |

**EFT subgroups' longitudinal overlap**

| | | | |
|---|---|---|---|
| BL overlaps FU2 Group A | 37 | FU2 overlaps FU3 Group A | 39 |
| BL overlaps FU2 Group B | 101 | FU2 overlaps FU3 Group B | 89 |
| BL&FU2 overlap | 138 | FU2&FU3 overlap | 128 |
| BL Group A overlaps FU2 Group A | 7 | FU2 Group A overlaps FU3 Group A | 34 |
| BL Group B overlaps FU2 Group B | 11 | FU2 Group B overlaps FU3 Group B | 68 |
| Consistent BL to FU2 Groups | 18 | Consistent FU2 to FU3 Groups | 102 |

(B)

**SST clustering results**

| | BL | FU2 | FU3 |
|---|---|---|---|
| Group A | 113 | 109 | 70 |
| Group B | 149 | 123 | 125 |

**SST subgroups' longitudinal overlap**

| | | | |
|---|---|---|---|
| BL overlaps FU2 Group A | 95 | FU2 overlaps FU3 Group A | 62 |
| BL overlaps FU2 Group B | 113 | FU2 overlaps FU3 Group B | 101 |
| BL&FU2 overlap | 208 | FU2&FU3 overlap | 163 |
| BL Group A overlaps FU2 Group A | 67 | FU2 Group A overlaps FU3 Group A | 43 |
| BL Group B overlaps FU2 Group B | 79 | FU2 Group B overlaps FU3 Group B | 51 |
| Consistent BL to FU2 Groups | 146 | Consistent FU2 to FU3 Groups | 94 |

**Table S3 EFT Subgroup Differences (Group A vs Group B) in measures of affective go/no-go task (AGNG) and the negative bias in Group B**

Group A featured high engagement with the dorsal anterior cingulate cortex (ACC) and the primary visual cortex (V1). Group B featured high engagement with the left inferior frontal cortex (L-IFC). Abbreviations: EFT, emotional face task; BL, baseline; FU2, follow-up 2; FU3, follow-up 3; n_Eff, effective sample size; fdr, false discovery rate.

**Experiment 1: Between-Group Difference in All AGNG Measures**

| EFT | Positive stimuli latency difference |
|---|---|
| BL Group A 120 vs Group B 179<br>n_Eff = 287.3579 | $t$=1.20, $p$=0.23, Cohen's $d$=0.14<br>$\boldsymbol{p_{fdr}}$ **= 0.41** |
| FU2 Group A 38 vs Group B 182<br>n_Eff = 125.7455 | $t$=0.89, $p$=0.38, Cohen's $d$ =0.16<br>$\boldsymbol{p_{fdr}}$ **= 0.57** |
| FU3 Group A 100 vs Group B 95<br>n_Eff = 194.8718 | $t$=1.45, $p$=0.15, Cohen's $d$ =0.21<br>$\boldsymbol{p_{fdr}}$ **= 0.36** |

| EFT | Negative stimuli latency difference |
|---|---|
| BL Group A 121vs Group B 180<br>n_Eff = 289.4352 | $t$=3.66, $p$=0.0030, Cohen's $d$=0.43<br>$\boldsymbol{p_{fdr}}$ **= 0.018** |
| FU2 Group A 39 vs Group B 181<br>n_Eff = 128.3455 | $t$=3.47, $p$=0.00070, Cohen's $d$=0.61<br>$\boldsymbol{p_{fdr}}$ **= 0.0084** |
| FU3 Group A 100 vs Group B 95<br>n_Eff = 194.8718 | $t$=2.84, $p$=0.0050, Cohen's $d$=0.41<br>$\boldsymbol{p_{fdr}}$ **= 0.020** |

| EFT | Positive stimuli omission difference |
|---|---|
| BL Group A 120 vs Group B 179<br>n_Eff = 287.3579 | $t$=-0.32, $p$=0.75, Cohen's $d$=-0.040<br>$\boldsymbol{p_{fdr}}$ **= 0.81** |
| FU2 Group A 38 vs Group B 182<br>n_Eff = 125.7455 | $t$=0.35, $p$=0.74, Cohen's $d$=0.070<br>$\boldsymbol{p_{fdr}}$ **= 0.81** |
| FU3 Group A 100 vs Group B 95<br>n_Eff = 194.8718 | $t$=-1.18, $p$=0.24, Cohen's $d$ =-0.17<br>$\boldsymbol{p_{fdr}}$ **= 0.41** |

| EFT | Negative stimuli omission difference |
|---|---|
| BL Group A 121vs Group B 180<br>n_Eff = 289.4352 | $t$=-0.27, $p$=0.79, Cohen's $d$=-0.031<br>$\boldsymbol{p_{fdr}}$ **= 0.81** |
| FU2 Group A 39 vs Group B 181<br>n_Eff = 128.3455 | $t$=-0.24, $p$=0.81, Cohen's $d$=-0.043<br>$\boldsymbol{p_{fdr}}$ **= 0.81** |
| FU3 Group A 100 vs Group B 95<br>n_Eff = 194.8718 | $t$=1.47, $p$=0.14, Cohen's $d$ =0.21<br>$\boldsymbol{p_{fdr}}$ **= 0.36** |

**Experiment 2: Within Group B Negative Bias in All AGNG Measures**

| EFT | L-IFC (GroupB) group positive stimuli latency vs negative stimuli latency |
|---|---|
| BL 179 | $t$=3.89, $p$=0.00014, Cohen's $d$=0.29<br>$p_{fdr}$ **= 0.00081** |
| FU2 181 | $t$=3.72, $p$=0.00027, Cohen's $d$=0.28<br>$p_{fdr}$ **= 0.00081** |
| FU3 95 | $t$=1.37, $p$=0.17, Cohen's $d$=0.14<br>$p_{fdr}$ **= 0.26** |

| EFT | L-IFC (GroupB) group positive stimuli omissions vs negative stimuli omissions |
|---|---|
| BL 179 | $t$=-0.046, $p$=0.65, Cohen's $d$< 0.010<br>$p_{fdr}$ **= 0.65** |
| FU2 181 | $t$=0.49, $p$=0.62, Cohen's $d$< 0.010<br>$p_{fdr}$ **= 0.65** |
| FU3 95 | $t$=-2.54, $p$=0.013, Cohen's $d$=-0.26<br>$p_{fdr}$ **= 0.026** |

**Experiment 3: Between-Group Emotional Symptoms and Depression Scores**

| EFT | Emotional Symptoms |
|---|---|
| BL Group A 122vs Group B 180<br>n_Eff = 290.8609 | $t$=-2.68, $p$=0.0078, Cohen's $d$ = -0.31<br>$p_{fdr}$ **= 0.012** |
| FU2 Group A 38 vs Group B 179<br>n_Eff = 125.3825 | $t$=-3.14, $p$=0.0025, Cohen's $d$ = -0.56<br>$p_{fdr}$ **= 0.0075** |
| FU3 Group A 99 vs Group B 93<br>n_Eff = 191.8125 | $t$=-2.67, $p$=0.0083, Cohen's $d$ = -0.39<br>$p_{fdr}$ **= 0.012** |

| EFT | Depression Score |
|---|---|
| BL Group A 121vs Group B 179<br>n_Eff = 288.7867 | $t$=1.85, $p$=0.065, Cohen's $d$=0.22<br>$p_{fdr}$ **= 0.065** |
| FU2 Group A 39 vs Group B 180<br>n_Eff = 128.2192 | $t$=-2.00, $p$=0.060, Cohen's $d$=-0.35<br>$p_{fdr}$ **= 0.065** |
| FU3 Group A 100vs Group B 95<br>n_Eff = 194.8718 | $t$=-3.75, $p$=0.00023, Cohen's $d$=-0.54<br>$p_{fdr}$ **= 0.0014** |

**Experiment 4: Persistent Low vs High Groups for Depression Scores**

| EFT | Emotional Symptoms |
|---|---|
| BL-FU2 Persist Group A 7 vs Persist Group B 11<br>n_Eff = 17.1111 | **BL**: $t$=-4.29, $p$=0.00049, Cohen's $d$=-2.07<br>$p_{fdr}$ **= 0.0020** |
| | **FU2** $t$=-2.81, $p$=0.012, Cohen's $d$=-1.35<br>$p_{fdr}$ **= 0.012** |
| FU2-FU3 Persist Group A 34 vs Persist Group B 68 | **FU2**: $t$=-2.75, $p$=0.0072, Cohen's $d$=-0.58 |

| | |
|---|---|
| n_Eff = 90.6667 | $p_{fdr}$ **= 0.012** |
| | **FU3** *t*=-2.65, *p*=0.0095, Cohen's *d* =-0.56<br>$p_{fdr}$ **= 0.012** |

**Table S4. Contingency tables of neurocognitive subgroups and “high-risk vs control” for MDD and AUD in IMAGEN at FU3.**

(A) In EFT, the left-IFC subgroup strongly predicted high-risk depression individuals (>50% chance of diagnosis according to DAWBA rating); (B) SST high pgACC/mPFC/SMA-engagement subgroup strongly predicted high risk in alcohol dependence (AUDIT total score >= 8). Abbreviations: DAWBA, the Development and Well-Being Assessment; AUDIT, the Alcohol Use Disorders Identification Test; IFC, inferior frontal cortex; dACC, dorsal anterior cingulate cortex; pgACC, pregenual anterior cingulate cortex; mPFC, medial prefrontal cortex; SMA, supplementary motor area.

(A)

| FU3 | High-Risk Depression | Controls |
|---|---|---|
| EFT Group B | 14 | 14 |
| EFT Group A | 3 | 27 |
| *OR* = 9.0 [2.2～36.6] | $\chi 2 = 11.2$ | $p = 8.3 \times 10^{-4}$ |

(B)

| FU3 | High-Risk Alcohol Dependence | Controls |
|---|---|---|
| SST Group B | 18 | 11 |
| SST Group A | 4 | 29 |
| *OR* = 11.9 [3.3~43.0] | $\chi 2 = 16.8$ | $p = 4.2 \times 10^{-5}$ |

**Table S5. Demographics between neurocognitive subgroups.**

No between-group difference was observed for emotional face task (EFT) or stop-signal task (SST) subgroups across baseline (BL), follow-up 2 (FU2), and follow-up 3 (FU3).

| EFT Group A vs B | gender | site |
|---|---|---|
| BL | $t$=-1.34, $p$=0.18, Cohen's $d$=-0.18 | $t$=-0.66, $p$=0.51, Cohen's $d$=-0.09 |
| FU2 | $t$=0.07, $p$=0.94, Cohen's $d$=-0.01 | $t$=-0.14, $p$=0.89, Cohen's $d$=-0.02 |
| FU3 | $t$=1.12, $p$=0.26, Cohen's $d$=0.15 | $t$=0.02, $p$=0.98, Cohen's $d$=0.00 |

| SST Group A vs B | gender | site |
|---|---|---|
| BL | $t$=0.22, $p$=0.82, Cohen's $d$=0.03 | $t$=0.39, $p$=0.69, Cohen's $d$=0.06 |
| FU2 | $t$=-1.03, $p$=0.30, Cohen's $d$=-0.14 | $t$=-0.28, $p$=0.77, Cohen's $d$=-0.04 |
| FU3 | $t$=-0.28, $p$=0.77, Cohen's $d$=-0.04 | $t$=-0.03, $p$=0.97, Cohen's $d$=-0.01 |

#### Table S6. SST Subgroup Differences (Group A vs Group B) in SST behaviors, AUDIT total scores and impulsivity measures

Abbreviations: SST, stop-signal task; AUDIT, the Alcohol Use Disorders Identification Test; BIS, the Barratt Impulsiveness Scale; CGT, the Cambridge Gambling Task; TCI-R, the Temperament and Character Inventory-Revised, SURPS, the Substance Use Risk Personality Scale.

**Experiment 1: Between-group difference in SST behaviors**

| SST | Stop reaction time difference |
|---|---|
| BL Group A 111 vs Group B 147<br>n_Eff = 252.9767 | $t$=1.30, $p$=0.19, Cohen's $d$=0.13<br>$\boldsymbol{p_{fdr}}$ **= 0.29** |
| FU2 Group 107 A vs Group B 121<br>n_Eff = 227.1404 | $t$=0.80, $p$=0.42, Cohen's $d$=0.08<br>$\boldsymbol{p_{fdr}}$ **= 0.42** |
| FU3 Group A 67 vs Group B 124<br>n_Eff = 173.9895 | $t$=1.05, $p$=0.29, Cohen's $d$=0.11<br>$\boldsymbol{p_{fdr}}$ **= 0.35** |

| SST | GO reaction time difference |
|---|---|
| BL Group A 111 vs Group B 147<br>n_Eff = 252.9767 | $t$=2.72, $p$=0.0070, Cohen's $d$ = 0.34<br>$\boldsymbol{p_{fdr}}$ **= 0.015** |
| FU2 Group 107 A vs Group B 121<br>n_Eff = 227.1404 | $t$=3.44, $p$=0.00069, Cohen's $d$=0.46<br>$\boldsymbol{p_{fdr}}$ **= 0.0041** |
| FU3 Group A 67 vs Group B 124<br>n_Eff = 173.9895 | $t$=2.70, $p$=0.0076, Cohen's $d$=0.41<br>$\boldsymbol{p_{fdr}}$ **= 0.015** |

**Experiment 2: Between-group differences for AUDIT**

| SST | AUDIT total score |
|---|---|
| BL Group A 110 vs Group B 145<br>n_Eff = 250.1961 | $t$=-4.31, $p$=0.000023, Cohen's $d$=-0.54<br>$\boldsymbol{p_{fdr}}$ **= 0.000023** |
| FU2 Group A 108 vs Group B 121<br>n_Eff = 228.262 | $t$=-4.39, $p$=0.000017, Cohen's $d$=-0.58<br>$\boldsymbol{p_{fdr}}$ **= 0.000023** |
| FU3 Group A 68 vs Group B 122<br>n_Eff = 174.6526 | $t$=-4.37, $p$=0.000021, Cohen's $d$=-0.66<br>$\boldsymbol{p_{fdr}}$ **= 0.000023** |

**Experiment 3: Persistent low vs high groups for AUDIT total score**

| SST | **AUDIT total score** |
|---|---|
| BL-FU2 Persist Group A 67 vs Persist Group B 79<br>n_Eff = 145.0137 | **BL**: $t$=-2.74, $p$=0.0068, Cohen's $d$ = -0.46<br>$\boldsymbol{p_{fdr}}$ **= 0.013** |
| | **FU2** $t$=-2.52, $p$=0.013, Cohen's $d$ = -0.42<br>$\boldsymbol{p_{fdr}}$ **= 0.013** |
| FU2-FU3 Persist Group A 43 vs Persist Group B 51<br>n_Eff = 93.31915 | **FU2**: $t$=-2.54, $p$=0.013, Cohen's $d$ = -0.53<br>$\boldsymbol{p_{fdr}}$ **= 0.013** |
| | **FU3** $t$=-2.71, $p$=0.008, Cohen's $d$ = -0.56<br>$\boldsymbol{p_{fdr}}$ **= 0.013** |

#### BIS

| SST Group A vs B | Attention | Motor | Planning |
|---|---|---|---|
| Follow-up 2 | *t* = 0.17, *p* = 0.85, Cohen's *d* = 0.02 | *t* = 1.31, *p* = 0.18, Cohen's *d* = 0.17 | *t* = 0.54, *p* = 0.58, Cohen's *d* = 0.07 |
| Follow-up 3 | *t* = 0.39, *p* = 0.69, Cohen's *d* = 0.06 | *t* = -0.55, *p* = 0.58, Cohen's *d* = 0.08 | *t* = 0.46, *p* = 0.64, Cohen's *d* = 0.06 |

#### CGT

| SST Group A vs B | Delay Aversion | Deliberation Time | Overall Proportion Bet | Quality of Decision-Making | Risk Adjustment | Risk Taking |
|---|---|---|---|---|---|---|
| Baseline | *t* = -0.50, *p* = 0.61, Cohen's *d* = -0.07 | *t*= 0.98, *p* = 0.32, Cohen's *d* = 0.15 | *t* = 0.84, *p* = 0.39, Cohen's *d* = 0.11 | *t* = 1.18, *p* = 0.23, Cohen's *d* = 0.16 | *t* = 0.20, *p* = 0.83, Cohen's *d* = 0.03 | *t* = 0.52, *p* = 0.6, Cohen's *d* = 0.07 |
| Follow-up 2 | *t* = 0.03, *p* = 0.97, Cohen's *d* = 0.00 | *t* = 0.07, *p* = 0.94, Cohen's *d* = 0.01 | *t* = -0.06, *p* = 0.95, Cohen's *d* = -0.01 | *t* = 0.09, *p* = 0.92, Cohen's *d* = 0.012 | *t* = -0.93, *p* = 0.34, Cohen's *d* = -0.12 | *t* = -0.17, *p* = 0.85, Cohen's *d* = 0.02 |
| Follow-up 3 | *t*= 0.17, *p* = 0.85, Cohen's *d* = 0.03 | *t* = -1.28, *p* = 0.2, Cohen's *d* = -0.18 | *t* = -0.77, *p* = 0.44, Cohen's *d* = -0.11 | *t* = 0.70, *p* = 0.48, Cohen's *d* = 0.10 | *t* = -0.22, *p* = 0.82, Cohen's *d* = -0.03 | *t* = -0.88, *p* = 0.37, Cohen's *d* = -0.13 |

#### Impulsivity Measures from SURPS and TCI-R

| SST Group A vs B | imp_mean (SURPS) | tci_imp (TCI-R) |
|---|---|---|
| Baseline | *t* = -0.45, *p* = 0.64, Cohen's *d* = -0.04 | *t* = 0.34, *p* = 0.73, Cohen's *d* = 0.03 |
| Follow-up 2 | *t* = 0.78, *p* = 0.43, Cohen's *d* = 0.08 | *t* = -0.67, *p* = 0.49, Cohen's *d* = -0.07 |
| Follow-up 3 | *t* = -0.32, *p* = 0.74, Cohen's *d* = -0.03 | *t* = 0.11, *p* = 0.91, Cohen's *d* = 0.01 |

#### Table S7. Hierarchical clustering results in the clinical Stratify cohort.

(A) Dendrograms for the emotion face task (EFT; left) in MDD+HC cohort and stop-signal task (SST; right) in AUD+HC cohort. (B) Silhouette coefficients suggest two-cluster solutions optimal (highlighted in bold red) for both EFT and SST in the respective cohorts.

##### (A) Dendrograms and clustering results

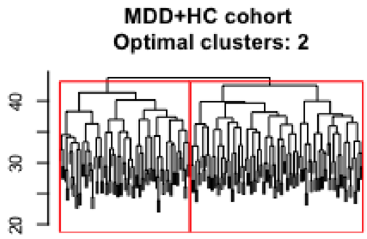


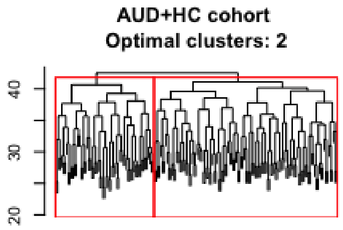


| | Group A | Group B |
|---|---|---|
| EFT | 77 | 116 |
| SST | 65 | 115 |

##### (B) Silhouette coefficients of clustering under different clustering numbers

| | 2 | 3 | 4 | 5 | 6 |
|---|---|---|---|---|---|
| EFT | **0.58** | 0.36 | 0.24 | 0.17 | 0.13 |
| SST | **0.61** | 0.42 | 0.37 | 0.33 | 0.27 |

#### Table S8. Diagnostic specificity of Brain-DiT models via counterfactual validation.

(A)-(B) Counterfactual analyses with EFT on alcohol use disorder (AUD) and SST on major depression disorder (MDD) confirmed the diagnostic specificity of Brain-DiT models. The nominal significance likely reflects comorbidities between AUD and MDD.

(A)

| | AUD | Control |
|---|---|---|
| EFT Group B | 70 | 26 |
| EFT Group A | 49 | 35 |
| $OR$ = 1.92 [1.03 ~ 3.59] | $\chi^2 = 4.25$ | $p = 0.039$ |

(B)

| | MDD | Control |
|---|---|---|
| SST Group B | 80 | 31 |
| SST Group A | 47 | 35 |
| $OR$ = 1.92 [1.05 ~ 3.51] | $\chi^2 = 4.56$ | $p = 0.033$ |

#### Table S9. Prediction Accuracy of MDD and AUD in Stratify with Machine Learning Classifiers based on activation, rs-FC, and CRS.

Common supervised machine learning (ML) approaches, Logistic regression, XGBoost, and SVM, were employed to train a prediction model from the IMAGEN cohort and predict MDD and AUD patients from controls in the Stratify cohort. Unsupervised method K-means (with cluster = 2) was also tested, with predefined input features (i.e., the key features identified by the main results: L-IFC vs dACC/V1 for the EFT and pgACC/mPFC/SMA for the SST). As a comparison, the unsupervised hierarchical clustering in the Stratify cohort with the whole brain CRS reached a classification accuracy of 85.0% for MDD vs controls and 82.2% for AUD vs controls. Abbreviations: MDD, the major depression disorder; AUD, the alcohol use disorder; rs-FC, resting-state functional connectivity; CRS, cognitive relevance score; EFT, emotional face task; IFC, inferior frontal cortex; dACC, dorsal anterior cingulate cortex; SST, stop-signal task; pgACC, pregenual anterior cingulate cortex; mPFC, medial prefrontal cortex; SMA, supplementary motor area.

(A) MDD with EFT

| Classifier/Clustering method | Modalities and Features from the Training Set IMAGEN | Classification Accuracy in the validation set STRATIFY |
|---|---|---|
| Logistic regression | Activation (key regions) | 0.53 |
| XGBoost | Activation (key regions) | 0.60 |
| SVM | Activation (key regions) | 0.59 |
| K-means | Activation (key regions) | 0.58 |
| Logistic regression | rs-FC | 0.68 |
| XGBoost | rs-FC | 0.65 |
| SVM | rs-FC | 0.68 |
| K-means | rs-FC | 0.58 |
| Logistic regression | CRS (key regions) | 0.50 |
| XGBoost | CRS (key regions) | 0.47 |
| SVM | CRS (key regions) | 0.64 |
| K-means | CRS (key regions) | 0.70 |

(B) AUD with SST

| Classifier/Clustering method | Modalities and Features from the Training Set IMAGEN | Classification Accuracy in the validation set STRATIFY |
|---|---|---|
| Logistic regression | Activation (key regions) | 0.54 |
| XGBoost | Activation (key regions) | 0.66 |
| SVM | Activation (key regions) | 0.66 |
| K-means | Activation (key regions) | 0.63 |
| Logistic regression | rs-FC | 0.64 |
| XGBoost | rs-FC | 0.63 |
| SVM | rs-FC | 0.62 |
| K-means | rs-FC | 0.56 |
| Logistic regression | CRS (key regions) | 0.64 |
| XGBoost | CRS (key regions) | 0.67 |
| SVM | CRS (key regions) | 0.62 |
| K-means | CRS (key regions) | 0.67 |

**Table S10. CRS (cognitive relevance score) features cannot be linearly derived from rs-FC and activation.**

**Target activation + rs-FC**

| **Task** | **Dataset** | **Average prediction accuracy (mean correlation of HCP-ex regions)** |
|---|---|---|
| Emotion face task | IMAGEN | Average $r$ = -0.01 (average $p$ = 0.48, all region $p$ > 0.05) |
| Emotion face task | Stratify | Average $r$ = 0.00 (average $p$ = 0.48, all region $p$ > 0.05) |
| Stop-signal task | IMAGEN | Average $r$ = -0.02 (average $p$ = 0.52, all region $p$ > 0.05) |
| Stop-signal task | Stratify | Average $r$ = 0.05 (average $p$ = 0.55, all region $p$ > 0.05) |

**Target activation + Input activation + rs-FC**

| **Task** | **Dataset** | **Average prediction accuracy (mean correlation of HCP-ex regions)** |
|---|---|---|
| Emotion face task | IMAGEN | Average $r$ = -0.01 (average $p$ = 0.41, all region $p$ > 0.05) |
| Emotion face task | Stratify | Average $r$ = 0.00 (average $p$ = 0.37, all region $p$ > 0.05) |
| Stop-signal task | IMAGEN | Average $r$ = -0.02 (average $p$ = 0.64, all region $p$ > 0.05) |
| Stop-signal task | Stratify | Average $r$ = 0.05 (average $p$ = 0.73, all region $p$ > 0.05) |

**Table S11. Brain-DiT architecture and hyperparameters.**

A. Embed part of Brain-DiT

| Layer | Filter size, Stride | Output size (C, D, H, W) |
|---|---|---|
| Input | - | 1x64x64x64 |
| Embed Layer | 8x8x8,8 | 8x8x8 |

B. Attention block part of Brain-DiT

| Parameter | Model value |
|---|---|
| Input dimension (dim) | [redacted] |
| Number of heads | [redacted] |
| MLP ratio | [redacted] |
| QKV bias | [redacted] |
| Activation function | [redacted] |
| Normalization layer | [redacted] |
| Rotary embed | [redacted] |
| Down-sampling block num | [redacted] |
| Middle block num | [redacted] |
| Up-sampling block num | [redacted] |

C. Hyperparameter of Brain-DiT

| Parameter | Value |
|---|---|
| Batch size | [redacted] |
| Nepoch | [redacted] |
| Embed dim | [redacted] |
| Patch size | [redacted] |
| Pred mode | [redacted] |
| Num head | [redacted] |
| P_uncond | [redacted] |
| Inference nstep | [redacted] |

(A) Embedding layer: 8x8x8 patch division → 512D features; (B) Attention: 8-head mechanism + RoPE positional encoding; (C) Training: batch 64/400 epochs/noise prediction mode, 50-step inference denoising. Total parameters: 75M (6.5 hours of training on a single A100 GPU).
